# Vimentin cysteine 328 modifications finely tune network organization and influence actin remodeling under oxidative and electrophilic stress

**DOI:** 10.1101/2023.03.30.534617

**Authors:** Patricia González-Jiménez, Sofia Duarte, Alma E. Martínez, Elena Navarro-Carrasco, Vasiliki Lalioti, María A. Pajares, Dolores Pérez-Sala

**Author notes:** To whom correspondence should be addressed at: Centro de Investigaciones Biológicas Margarita Salas, C.S.I.C., Ramiro de Maeztu, 9, 28040 Madrid, Spain.

## Abstract

Cysteine residues can undergo multiple posttranslational modifications with diverse functional consequences, potentially behaving as tunable sensors. The intermediate filament protein vimentin has important implications in pathophysiology, including cancer progression, infection, and fibrosis, and maintains a close interplay with other cytoskeletal structures, such as actin filaments and microtubules. We previously showed that the single vimentin cysteine, C328, is a key target for oxidants and electrophiles. Here, we demonstrate that structurally diverse cysteine-reactive agents, including electrophilic mediators, oxidants and drug-related compounds, disrupt the vimentin network eliciting morphologically distinct reorganizations. As most of these agents display broad reactivity, we pinpointed the importance of C328 by confirming that local perturbations introduced through mutagenesis provoke structure-dependent vimentin rearrangements. Thus, GFP-vimentin wild type (wt) forms squiggles and short filaments in vimentin-deficient cells, the C328F, C328W, and C328H mutants generate diverse filamentous assemblies, and the C328A and C328D constructs fail to elongate yielding dots. Remarkably, vimentin C328H structures resemble the wt, but are strongly resistant to electrophile-elicited disruption. Therefore, the C328H mutant allows elucidating whether cysteine-dependent vimentin reorganization influences other cellular responses to reactive agents. Electrophiles such as 1,4-dinitro-1H-imidazole and 4-hydroxynonenal induce robust actin stress fibers in cells expressing vimentin wt. Strikingly, under these conditions, vimentin C328H expression blunts electrophile-elicited stress fiber formation, apparently acting upstream of RhoA. Analysis of additional vimentin C328 mutants shows that electrophile-sensitive and assembly-defective vimentin variants permit induction of stress fibers by reactive species, whereas electrophile-resistant filamentous vimentin structures prevent it. Together, our results suggest that vimentin acts as a break for actin stress fibers formation, which would be released by C328-aided disruption, thus allowing full actin remodeling in response to oxidants and electrophiles. These observations postulate C328 as a “sensor” transducing structurally diverse modifications into fine-tuned vimentin network rearrangements, and a gatekeeper for certain electrophiles in the interplay with actin.

## 1. Introduction

Intermediate filaments of the type III family, which comprises vimentin, glial fibrillary acidic protein, peripherin and desmin, play key roles in the integration of cellular functions and cytoskeletal responses to stress. Proteins of this family are expressed in a cell-type specific manner and form extended cytoplasmic filament networks interplaying with tubulin and actin [1, 2]. The key role of these proteins in cell homeostasis is highlighted by the severe diseases arising from mutations in their sequence. These pathologies span from cataract and premature aging syndrome, as reported for vimentin mutations [3, 4], to fatal neurodegenerative disease in the case of the glial fibrillary acidic protein (GFAP) [5], and skeletal and cardiac myopathies caused by desmin mutations [6]. Vimentin is perhaps the most thoroughly studied member of this family, both in biological systems and *in vitro*. Vimentin is present both in the cytoplasm and in the extracellular medium and plays important roles in cell migration, division, regulation of the inflammatory response, interaction with pathogens, tumorigenic transformation, etc. [7–11].

Therefore, type III intermediate filaments, and particularly vimentin, are considered important drug targets in proliferative, autoimmune and infectious diseases [10, 12, 13]. Interestingly, studies in vimentin knockout animals or deficient cells have unveiled that many vimentin functions become especially obvious upon injury or stress [14]. Indeed, compromised wound healing, impaired regulation of vascular tone after renal injury and deregulated inflammatory responses are some of the consequences of vimentin deficiency [14]. Importantly, oxidative stress is a hallmark of numerous pathophysiological processes including inflammation, ischemia reperfusion injury, neurological and cardiovascular disease, and cancer [15]. Interestingly, reactive oxygen, nitrogen and electrophilic species have been shown to deeply alter vimentin organization [16–18], both through direct modification of the protein, and indirectly through modulation of signaling pathways [12, 19].

Structurally, the vimentin monomer comprises an α-helical rod domain flanked by intrinsically disordered head and tail domains. Vimentin is believed to immediately form parallel dimers, which then assemble in antiparallel tetramers. The current models of filament assembly propose the formation of unit length filaments (ULF) of approximately eight tetramers associated side by side. These units would then imbricate end-to end to form filaments, which subsequently compact radially [20]. These processes imply marked conformational changes, which affect multiple segments of the protein, including the head and tail domains [21, 22]. In turn, vimentin disassembly is accompanied by a loosening of these conformations, and increased accessibility of vimentin domains [23].

Importantly, type III intermediate filament proteins possess a cysteine residue that is conserved across species and that, except in peripherin, is the only cysteine residue of the protein. Accumulating evidence indicates that this single cysteine residue, C328 in vimentin, is a target for oxidative and electrophilic modifications in several experimental models, as well as in human disease, thus behaving as a hot spot for posttranslational modifications (PTMs) [12, 24–27]. This could be related to the high reactivity of this residue, which in vimentin exhibits a low pKa, and is involved in zinc binding [28]. Moreover, detailed experiments in cultured cells as well as *in vitro* using purified vimentin, indicate that modifications of C328 have important consequences for vimentin assembly and filament morphology [18, 29], supporting a potential role for this residue in redox and electrophile/stress sensing [16, 30]. Interestingly, it appears that structurally diverse cysteine modifications could affect vimentin assembly in different ways [12, 18, 31]. Several electrophiles, including the endogenous mediators 15-deoxy-Δ^12,14^-prostaglandin J_2_ (15d-PGJ_2_) and 4-hydroxynonenal (HNE), or oxidants such as diamide or oxidized glutathione, limit filament elongation or provoke protein aggregation *in vitro*, while inducing filament condensation or fragmentation in cells [16, 18]. Given the key role of vimentin in cellular functions in health and disease, establishing the structure-function relationships of cysteine modifications could help understand its regulation and potentially aid to design therapeutic strategies.

Here we have explored the impact of several cysteine-reactive small electrophilic compounds, including endogenous mediators and drug analogs (Fig. 1A), on the distribution of vimentin constructs in vimentin-deficient cells. Moreover, the importance of structural changes at position 328 in vimentin assembly and in the response to stress has been evaluated by the use of vimentin mutants. Our results unveil the responsiveness of the C328 site to structural changes that, not only are translated into distinct vimentin network rearrangements, but also modulate the response of the cellular actin cytoskeleton to various electrophiles. Thus, based on evidence of C328 modifications by diverse reactive species [24, 29, 31–35], and the features highlighted herein, namely, the ability of changes at this site to elicit vimentin remodeling and to influence other cellular responses, a role for C328 as a sensor for oxidative and electrophilic stress is proposed.

**Figure 1.**
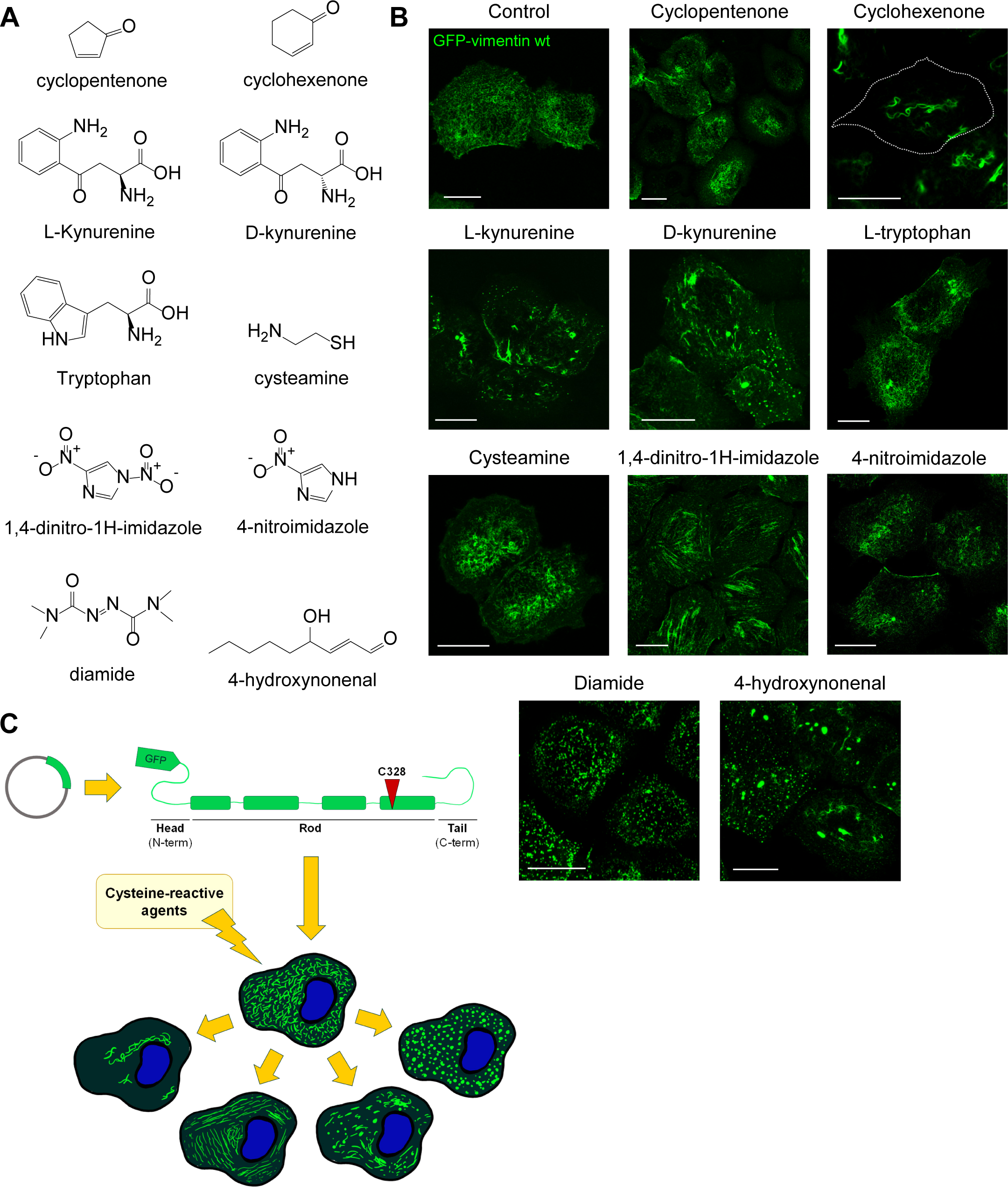
Reorganization of GFP-vimentin wt structures upon treatment of SW13/cl.2 cells with several cysteine-reactive agents. (A) Structure of various compounds used in this study. (B) SW13/cl.2 vimentin-deficient cells stably transfected with GFP-vimentin wt were treated with the indicated electrophilic compounds, under the conditions specified in the experimental section. After treatment, cells were fixed and the distribution of GFP-vimentin wt was monitored by fluorescence microscopy. Images shown are representative overlay projections from at least three assays per experimental condition. (C) Schematic view of the organization of a GFP-vimentin wt monomer (above), depicting the N-terminal (N-term) head, C-terminal (C-term) tail, and central (Rod) domains, highlighting the position of the single vimentin cysteine residue (C328). The cartoon below shows a representation of the typical squiggles formed by GFP-vimentin wt in vimentin-deficient cells (center), together main morphological reorganization patterns arising upon treatment with the cysteine-reactive agents shown in (A). In contrast to the homogeneous lattice of short filaments characteristic of the control condition, the various compounds elicited the rearrangement of short filaments into, from left to right, perinuclear condensations or filament retraction from the cell periphery, parallel linear arrays, bundles and accumulations, or bright dots. In some cases, mixed patterns could be observed. Bars, 20 μm.

## 2. Materials and Methods

### Materials

4-Hydroxynonenal (HNE) was from Cayman Chemical. L- and D-Kynurenine, cysteamine, L-tryptophan, 2-cyclopentenone, 2-cyclohexenone, 1,4-dinitro-1H-imidazole (DNI), 4-nitroimidazole (NI), and diamide were from Sigma. Oligonucleotides for mutagenesis were synthesized by Integrated DNA Technologies. Anti-vimentin, clones V9 (sc-6260) and E5 (sc-373717) and their Alexa fluorescent conjugates were from Santa Cruz Biotechnology; horseradish peroxidase (HRP)-conjugated secondary antibodies were from DAKO. Y27632 was from Calbiochem; calpeptin was from Cytoskeleton. (+)-Biotin-(PEO)3-iodoacetamide (biotinylated iodoacetamide) was from Santa Cruz Biotechnology. EZ-Link™ Maleimide-PEG2-Biotin (biotinylated maleimide) was from ThermoFisher Scientific.

### Cell culture and treatments

SW13/cl.2 vimentin-deficient cells from adrenal carcinoma were the generous gift of Dr. A. Sarriá (University of Zaragoza, Spain), and have been previously described [8, 16]. Murine embryonic fibroblasts (MEF) expressing endogenous vimentin were the generous gift of Prof. J. E. Eriksson (Abo Academy, Finland). Human breast carcinoma MCF7 (HTB-22™) cells were originally from ATCC and were authenticated by microsatellite amplification (short tandem repeat (STR)-PCR profiling) at Secugen, S.L. (Madrid, Spain). All cells were cultured in DMEM with 4.5 g/l glucose and glutamine (Invitrogen), plus 10% (v/v) fetal bovine serum (FBS; Sigma or Biowest) and antibiotics (100 U/ml penicillin and 100 μg/ml streptomycin; Invitrogen). All cells were routinely tested for the presence of Mycoplasma and confirmed to be free of contamination. Unless otherwise specified, treatment with the various agents was carried out in serum-free medium at 37°C. Whenever dimethylsulfoxide (DMSO) was used as solvent, control cells were incubated with an equivalent volume of this vehicle. Cyclopentenone and cyclohexenone were added at 100 µM for 3 h; L-, D-kynurenine and L-tryptophan were added at 1 mM for 3 h; cysteamine was used at 250 µM for 2 h; DNI and NI were added at 15 µM for 2 h. HNE was employed at 10 µM for 4 h; diamide was added at 1 mM for 15 min, and H_2_O_2_ was used at 1 mM for 30 min. The RhoK inhibitor Y27632, was used at 10 µM for 30 min, and calpeptin was used at 270 µM (1 ud/mL) for 20 min.

### Plasmids and transfections

The plasmids coding for fusion proteins of human vimentin and the green fluorescent protein, GFP-vimentin wt and the C328S and C328A mutants, have been previously described [16]. Additional mutants, GFP-vimentin C328F, C328W, C328H and C328D were generated by site directed mutagenesis using kits from Agilent or Nzytech, and the oligonucleotides depicted in Suppl. Table 1, following the instructions of the respective manufacturers. The bicistronic plasmid RFP//vimentin wt, coding for vimentin and the red fluorescent protein (DsRed Express2, RFP) as separate products was described in [16]. The RFP//vimentin C328H plasmid was generated by mutagenesis as above using the corresponding primers (Suppl. Table 1). The pRK5-RhoA L63Q plasmid, encoding a constitutively active RhoA was described in [36]. Cells were transfected with the indicated plasmids routinely using 1 μg of DNA mixed with 3 μl of Lipofectamine 2000 (Invitrogen) in Optimem (Gibco), per p35 dish. Experiments with transiently transfected cells were performed 48 h after transfection. Stable transfectants were obtained by selection in medium containing the antibiotic G-418 (500 μg/ml, final concentration), for at least 5 passages.

### Cell lysis and western blot

Cells in p60 dishes were washed twice with cold PBS on ice and the cell monolayer was lysed in 50 mM Tris–HCl pH 7.5, 0.1 mM EDTA, 0.1 mM EGTA, 0.1 mM β-mercaptoethanol, containing 0.5% (w/v) SDS, 0.1 mM sodium orthovanadate and protease inhibitors (2 μg/ml each of leupeptin, aprotinin and trypsin inhibitor, and 1.3 mM Pefablock), by gentle scraping and forced passes through a 26 1/2G needle. Cell debris was removed by centrifugation at 20,000 g for 5 min at 6°C. Protein concentration in cleared lysates was determined by the bicinchoninic acid (BCA) method (Thermo). Lysates were denatured at 95°C for 5 min in Laemmli buffer and aliquots containing 30 μg of protein were separated on 10 % (w/v) SDS-polyacrylamide gels. Proteins were transferred to Immobilon-P membranes (Millipore) using a semi-dry transfer system from Bio-Rad. Blots were blocked with 2% (w/v) powdered skimmed milk in T-TBS (20 mM Tris-HCl, pH 7.5, 500 mM NaCl, 0.05% (v/v) Tween-20), before incubation with primary antibodies, at 1:1000 dilution, followed by HRP-conjugated secondary antibodies at 1:2000 dilution, prepared in both cases in 1% (w/v) BSA in T-TBS. Bands of interest were visualized by chemiluminiscence using the ECL system (Cytiva), exposing blots to X-Ray film (Agfa).

### Immunoblot detection of protein carbonyl groups

Protein carbonyl formation by oxidative reactions was assessed with the OxyBlot Protein Oxidation Detection Kit (S7150, Merck Millipore), according to manufacturer’s instructions. Briefly, aliquots from cell lysates containing 30 µg of protein were denatured by addition of SDS at 6% (w/v) final concentration, carbonyl groups were derivatized 2,4-dinitrophenylhydrazine (DNPH) to obtain the corresponding 2,4-dinitrophenylhydrazones (DNP). Subsequently, the reaction mixtures were neutralized as instructed in the kit, and samples were then separated by SDS-PAGE and transferred to Immobilon-P membranes, as above. For dot blot, aliquots of derivatized lysates containing 15 µg of protein were spotted on a Whatman Protran nitrocellulose membrane (Perkin Elmer). Blots were blocked as above, before overnight incubation with the anti-dinitrophenyl antibody (anti-DNP) at 1:150 dilution, followed by overnight incubation with HRP-conjugated secondary antibody at 1:300 dilution. Results were visualized by ECL detection.

### Cysteine accessibility assay

SW13/cl.2 cells stably transfected with RFP//vim wt were incubated in control conditions or with DNI, and lysed as described above. Cellular debris was removed by centrifugation at 2,600 g for 5 min at 6°C. Lysates from control and DNI treated cells were split into two aliquots, and DTT was added to one of them at 1 mM, final concentration. All aliquots were then incubated on ice for 30 min, after which, 10 mM biotinylated iodoacetamide or biotinylated maleimide were added for free thiol group labeling, and incubation was continued for 30 min at room temperature. Samples were then used for immunoprecipitation.

### Immunoprecipitation

Agarose-conjugated anti-vimentin V9 beads (Santa Cruz Biotechnology sc-6260) were equilibrated in binding buffer containing 50 mM Tris–HCl pH 7.5, 0.1 mM EDTA, 0.1 mM EGTA, with 0.1% (w/v) SDS, 0.8% (v/v) NP-40, containing sodium orthovanadate and protease inhibitors as specified earlier. Cell lysates labeled with biotinylated iodoacetamide as above, were incubated with beads in binding buffer in a rotating wheel, for 1 h at 4°C. Beads were washed three times with cold PBS by centrifugation at 14,000 g for 1 min at 4°C, and subsequently boiled in Laemmli buffer during 5 min to elute the retained proteins. The immunoprecipitated products were separated and analyzed by SDS-PAGE and western blot.

### Fluorescence microscopy

The distribution of GFP-vimentin was assessed by confocal fluorescence microscopy. For fixation, cells were washed twice with PBS and incubated in 4% (w/v) paraformaldehyde (PFA) in PBS for 25 min at room temperature. For immunofluorescence detection of vimentin, fixed cells were permeabilized with 0.1% (v/v) Triton X-100 in PBS for 20 min at room temperature, blocked by incubation with 1% (w/v) BSA in PBS for 1 h, and incubated with Alexa488, Alexa546 or Alexa647-conjugated anti-vimentin antibody at 1:200 dilution in blocking solution for 1 h. For f-actin detection, fixed and permeabilized cells were incubated with 0.25 µg/ml phalloidin-tetramethylrhodamine B isothiocyanate (phalloidin-TRITC) (Sigma) in blocking solution. Images were acquired on SP5 or SP8 Leica confocal microscopes, using 63x magnification, routinely every 0.5 μm. Settings on the confocal microscope were saved and used in subsequent experiments with minor adjustments. LUT command was used to ensure non-overexposed acquisition. Where indicated, the Lightning module of the SP8 microscope was used for improved resolution.

### Image analysis

For image analysis, Leica LasX software or Image J (FIJI) were used. Single sections of images or overlay projections are shown, as indicated. The proportion of cells with disrupted GFP-vimentin was obtained by visually counting cells with vimentin dots, accumulations and/or parallel linear arrangements of vimentin squiggles. The proportion of cells displaying stress fibers was obtained by visual inspection of phalloidin-stained preparations.

Cells with linear actin structures longer than 5 µm, as measured with the Leica LasX ruler, were considered positive. Fluorescence intensity profiles of single sections were obtained, and the coefficient of variation calculated as the ratio between the standard deviation and the average value of fluorescence intensity. F-actin anisotropy, assessed with the Image J plugin “FibrilTool”, was used as a measure of the parallel disposition of stress fibers [37]. The anisotropy values were measured in manually defined regions of interest (ROI) extending from the cell cortex to two thirds of the distance between the cell membrane and the nucleus.

### Statistical analysis

All experiments were performed at least three times and measurements from distinct samples were taken. Results are presented as average values ± standard error of mean (SEM). Statistical analysis was performed with GraphPad Prism 8.0 software. For comparisons of multiple data sets the one-way analysis of variance (ANOVA) followed by Tukey’s multiple comparison test was used. For comparisons between two groups of data the Student’s *t*-test for unpaired samples was used. Statistically significant differences are indicated on the graphs and/or in the figure legends, as follows: *p<0.05, **p<0.005, ***p<0.001, ****p<0.0001. The number of determinations for each experimental condition or the sample size are given in the corresponding figure legends.

## 3. Results

### Treatment of cells with several cysteine-reactive agents results in distinct remodeling of GFP-vimentin

Vimentin-deficient SW13/cl.2 cells, which also lack other cytoplasmic intermediate filaments, constitute an adequate model to study the behavior of vimentin constructs upon transfection [38, 39]. Here, SW13/cl.2 cells stably transfected with GFP-vimentin wt were used to study the impact of several cysteine-reactive agents on vimentin assembly. Compounds used were selected on the basis of their reported reactivities and their structural similarity to endogenous metabolites/mediators or to therapeutic agents (Fig. 1A). Thus, 2-cyclopentenone mimics the core reactive structure of the endogenous lipid mediators known as cyclopentenone prostaglandins [40]; 2-cyclohexenone is the reactive moiety of several anticancer compounds [41]; L-kynurenine is a product of tryptophan degradation by indoleamine 2,3-dioxygenase [42], and it is used here at the concentrations reached in experimental in vivo studies [43]; cysteamine is an endogenous aminothiol employed for the treatment of cystinosis [44]; 1,4-dinitro-1H-imidazole (DNI), and its mono-nitrated counterpart, 4-nitroimidazole (NI), bear analogy with nitroimidazole compounds used as antiparasitic agents, and previously reported to form adducts with parasite proteins [45]. The effects of these compounds on GFP-vimentin wt organization are illustrated in Fig. 1B. As shown previously [16, 46], GFP-vimentin wt fusion constructs are unable to produce long filaments in vimentin-deficient cells, but form a uniform lattice constituted by squiggles and short filaments (Fig. 1B). These constructs can undergo remodeling in response to various stimuli [8, 16]. Here, cyclopentenone-treated cells showed apparently more robust squiggles, together with some bundles and accumulations, whereas cyclohexenone treatment resulted in the appearance of curly bundles. In turn, both D- and L-enantiomers of kynurenine [47] induced the formation of longer filamentous structures, together with some wavy bundles and dots (Fig. 1B).

Supplementing the medium with an equivalent concentration of L-tryptophan elicited milder alterations, with the appearance of some accumulations or short bundles, but preservation of squiggles in most cells (Fig. 1B). Treatment with cysteamine did not induce appreciable changes in GFP-vimentin wt distribution (Fig. 1B). Remarkably, DNI, which can react with cysteine residues introducing an imidazole group [48], elicited a complex reorganization pattern with the condensation of GFP-vimentin squiggles into various structures that frequently adopted a linear disposition in parallel arrangements (Fig. 1B). In contrast, in NI-treated cells, some peripheral condensation of vimentin was apparent, but squiggles were more preserved, and parallel arrays were less pronounced (Fig. 1B). Comparison of DNI and NI structures suggests that the peculiar effect of DNI could be related to the presence of a second nitro moiety. Finally, the effects of diamide and HNE are shown for reference (Fig. 1B). As we previously reported [16] diamide elicited a severe reorganization of GFP-vimentin wt filaments into dots, whereas HNE, at concentrations in the range of those measured under pathological conditions [49], provoked a more heterogeneous remodeling into dots and accumulations (Fig. 1B).

Taken together, these results indicate that several agents with the capacity to form covalent adducts with proteins induce distinct morphological patterns of reorganization of GFP-vimentin wt constructs in cells. A schematic view of the various GFP-vimentin arrangements observed is depicted in Fig. 1C.

### Cysteine-reactive agents induce remodeling of the cellular vimentin network

Next, we explored the effect of some of the electrophilic agents on the organization of a full vimentin network (Fig. 2). As we previously showed [16], SW13/cl.2 stably transfected with RFP//vimentin wt, therefore expressing untagged vimentin, displayed a homogeneous filament network formed by filaments or bundles of moderate thickness, extending from the nucleus towards the cell periphery (Fig. 2A). Treatment with cyclohexenone induced the appearance of accumulations or condensations of filaments together with curly bundles, whereas L-kynurenine elicited milder bundling (Fig. 2A). In turn, treatment with DNI induced a distinct pattern reminiscent of that observed in GFP-vimentin wt-expressing cells, with arrangement of vimentin filaments in linear parallel arrays, condensations or bundles (Fig. 2A). As previously reported [16], HNE and diamide, shown for reference, elicited vimentin filament retraction from the cell periphery and fragmentation into dots, respectively (Fig. 2A).

**Figure 2.**
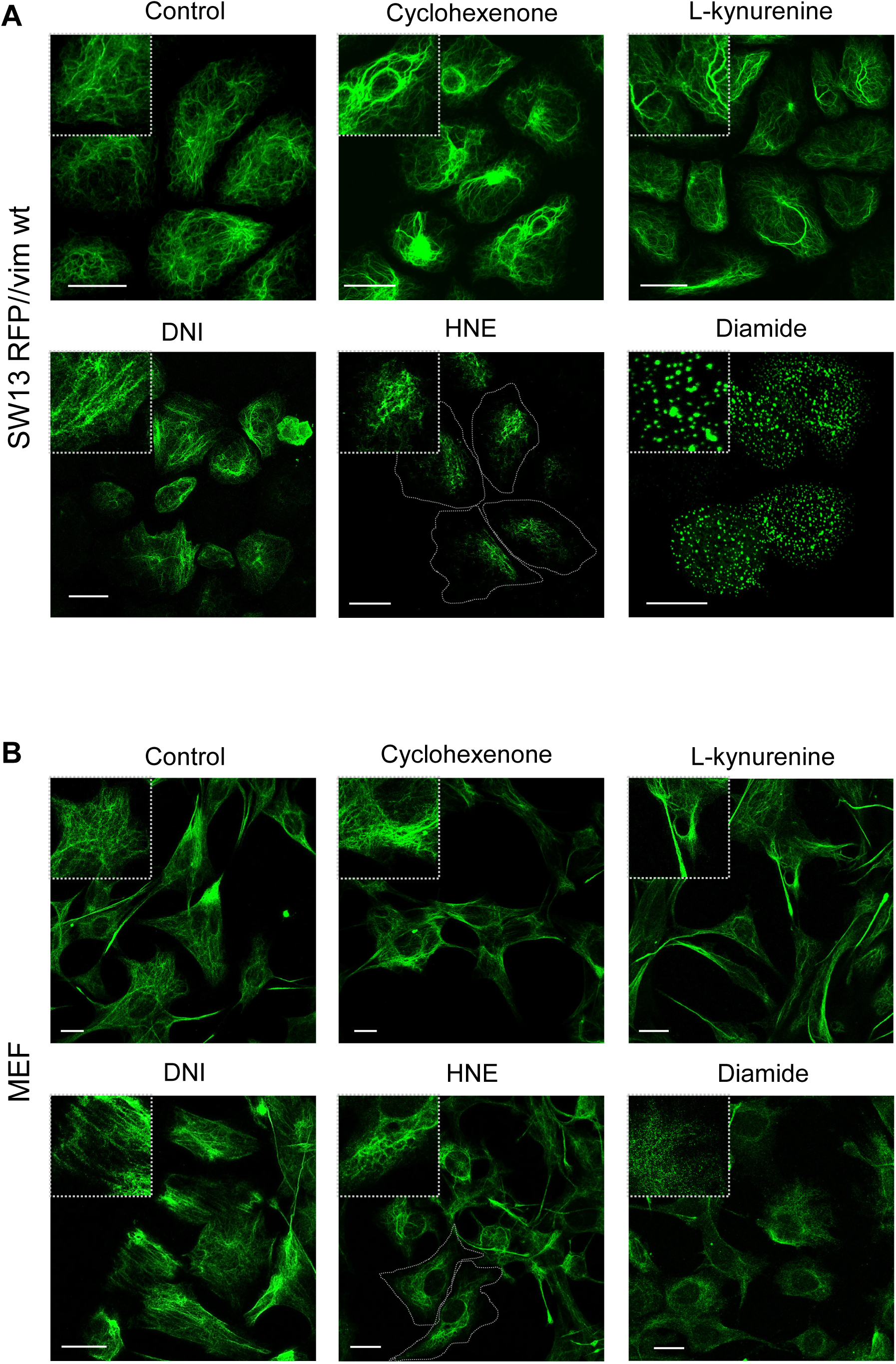
Effect of various electrophiles on the morphology of the vimentin network. SW13/cl.2 cells stably transfected with RFP//vimentin wt (A) or MEF expressing endogenous vimentin (B) were treated with the indicated electrophiles as in Fig. 1, and the morphology of the vimentin network was assessed by immunofluorescence. Images shown are overall projections from at least three assays per experimental condition. Insets show enlarged views of areas of interest. Cell contours are outlined in some images to illustrate vimentin network retraction from the cell periphery. For space reasons, in the labels of several figures, vimentin is abbreviated as “vim”. Bars, 20 μm.

Additionally, we explored the effect of the same electrophilic agents on the organization of endogenous vimentin in MEFs (Fig. 2B). Again, the results illustrate vimentin remodeling in morphologically diverse patterns depending on the agent used. Thus, MEFs treated with cyclohexenone or kynurenine displayed bundles and accumulations, whereas DNI elicited the frequent disposition of vimentin in linear parallel arrays, as observed in SW13/cl.2 cells. HNE-elicited filament retraction from the cell periphery, whereas diamide induced filament fragmentation, but yielded smaller dots than in SW13/cl.2 cells (Fig. 2B).

Taken together, our results show that various oxidants and electrophiles induce the remodeling of both endogenous and transfected (untagged or tagged) vimentin yielding different patterns, which are more prominent in the case of GFP-vimentin in relation with the assembly limitations imposed by the GFP tag, as we previously noted [16].

### Role of perturbations at position 328 of vimentin in network remodeling

Noteworthily, the electrophilic agents employed are broadly reactive and can potentially modify various nucleophilic residues, not only on vimentin but also on other cellular proteins. As a matter of fact, most electrophiles can form adducts with histidine and/or lysine residues, as reported for kynurenine [50, 51] and HNE [52]. Moreover, any single electrophile can provoke a chain reaction with generation of additional reactive species which prompt secondary modifications [53–55]. Indeed, we observed that treatment with several of the compounds led to an increase in the content of carbonyl groups in several cellular polypeptides, as detected using Oxyblot (Suppl. Fig. 1). Therefore, the reorganizations of the vimentin network observed above could be the result of both direct and indirect modifications. To assess the significance of perturbations specifically affecting the 328 site in vimentin remodeling, we adopted a strategy based on site-directed mutagenesis of GFP-vimentin, to yield the C328H, C328F and C328W substitutions, which introduce moieties weakly reminiscent of the adducts potentially formed by some of the reactive compounds used above. Next, we evaluated the impact of these mutations on vimentin network assembly and morphology. In SW13/cl.2 cells, assemblies formed by the various mutants showed evident differences with respect to GFP-vimentin wt organization (Fig. 3). The GFP-vimentin C328H mutant formed abundant short filaments which appeared thicker/more robust than the wt. GFP-vimentin C328F formed filamentous structures considerably longer than those of the wt, and frequently appearing as curly bundles. GFP-vimentin C328W squiggles were also longer and often distributed along the cell periphery, with some cells presenting intense bundles. The previously reported GFP-vimentin C328A and C328S mutants, unable to elongate and forming only bright dots [16], are shown for comparison (Fig. 3).

**Figure 3.**
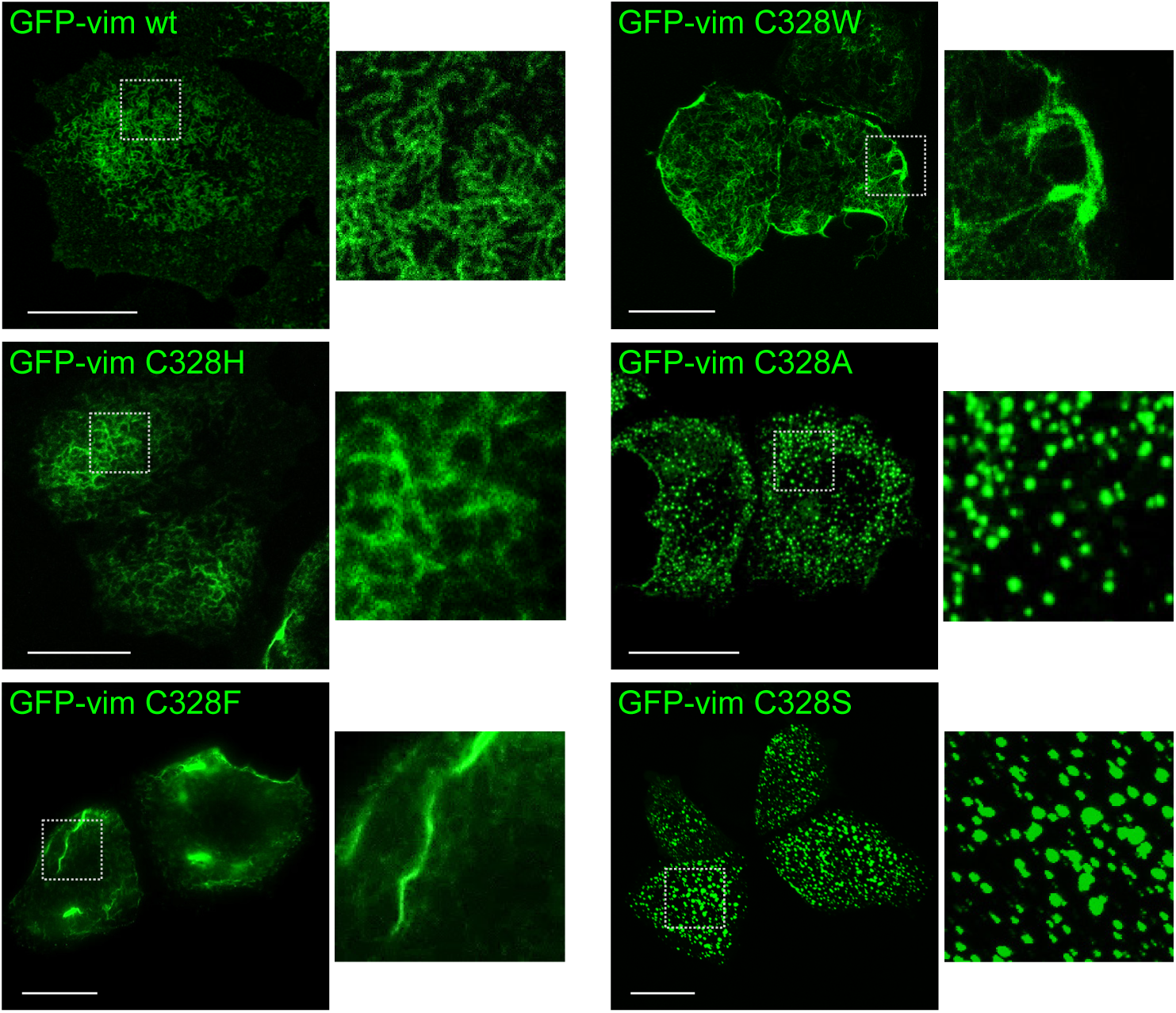
Morphology of the structures formed by GFP-vimentin wt and various cysteine mutants in vimentin-deficient cells. SW13/cl.2 were transfected with the indicated GFP fusion constructs of vimentin wt or cysteine mutants and the morphology of the assemblies formed was assessed 48 h later by fluorescence microscopy. Details of the assemblies formed by the various constructs are shown at the right of each image. Bars, 20 μm.

The assembly of GFP-vimentin cysteine mutants also showed striking differences in breast cancer MCF7 cells, which lack vimentin but express other cytoplasmic intermediate filaments, such as keratins [56] (Suppl. Fig. 2A). GFP-vimentin wt formed scattered thin filaments coexisting with abundant dots, and the GFP-vimentin C328S mutant formed only dots. In sharp contrast, GFP-vimentin C328H assembled into robust filamentous structures and bundles, whereas in cells expressing GFP-vimentin C328F or C328W, filaments and bundles coexisted with shorter wavy filaments. Analysis by western blot showed that the robustness of assemblies did not correlate with protein levels or alterations in size of the various mutant constructs (Suppl. Fig. 2B).

Therefore, in both SW13/cl.2 and MCF7 cells, the presence of a ring-like or aromatic moiety in the place of the cysteine residue appears to favor the formation of assemblies thicker and/or longer than those achieved by GFP-vimentin wt. Taken together, these observations show that point mutations at the 328 position differentially affect GFP-vimentin assembly.

### Vimentin C328H is more resistant than the wt protein to several oxidants and electrophiles

Cysteine residues are highly nucleophilic and prone to numerous oxidative modifications due to the presence of the sulfur center, whereas histidine residues possess lower nucleophilicity and are less susceptible to oxidation [57]. Given the ability of GFP-vimentin C328H to form robust short filaments in cells and its expected lower susceptibility to modification, we focused on this mutant to explore its response to various electrophiles and oxidants. As we previously reported [16] and illustrated above, treatment of cells expressing GFP-vimentin wt with the oxidant diamide or the electrophilic lipid HNE elicited a marked remodeling consisting in the transformation of vimentin squiggles into dots or accumulations, which occurred in virtually all cells treated with diamide and nearly 90% of those exposed to HNE (Fig. 4). Remarkably, GFP-vimentin C328H was practically resistant to the effect of diamide and largely resistant to that of HNE, as shown in Figure 4. Moreover, GFP-vimentin C328H structures were also resistant to treatment with DNI, since the typical squiggles conserved their wavy morphology and apparently uniform distribution, and did not rearrange into the parallel linear alignments adopted by GFP-vimentin wt upon DNI exposure. As expected, cells expressing GFP-vimentin C328H were also protected from the less potent effects of NI on GFP-vimentin wt remodeling, as well as from the disruption induced by H_2_O_2_ (30% vs 80% affected cells in GFP-vimentin C328H and wt, respectively). Indeed, quantitation of the proportion of cells undergoing morphological alterations of GFP-vimentin assemblies in response to the various agents confirmed the marked resistance of GFP-vimentin C328H to disruption under all conditions studied (Fig. 4A, graph).

**Figure 4.**
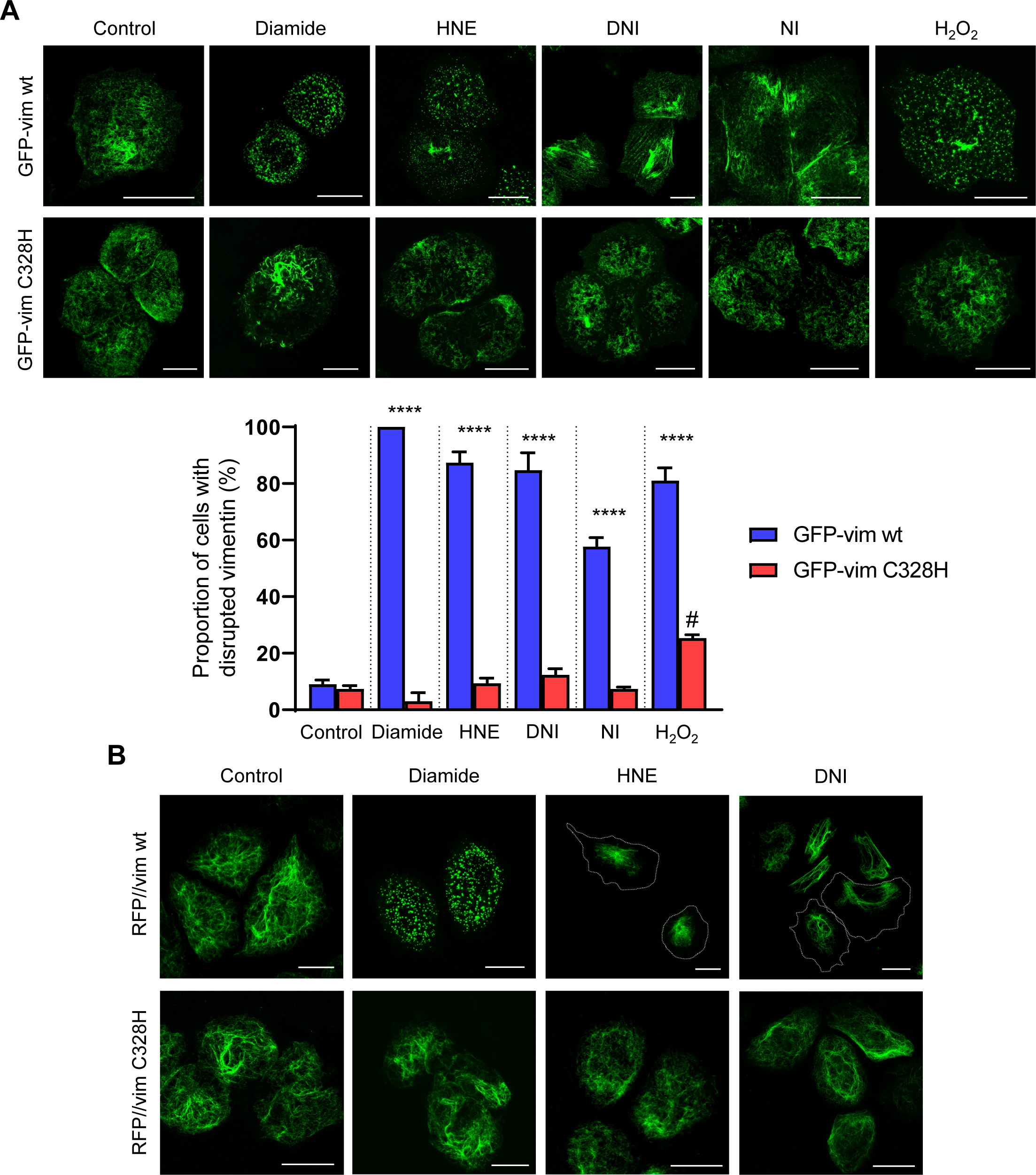
Effect of electrophiles and oxidants on the assemblies formed by vimentin wt or C328H. (A) SW13/cl.2 cells stably transfected with GFP-vimentin wt or C328H were treated with the indicated agents, as described in the experimental section. The morphology of vimentin assemblies was assessed by fluorescence microscopy. In the lower graph, the proportion of cells with disrupted vimentin, i.e., showing dots, accumulations and/or parallel arrays of GFP-vimentin was obtained by visual inspection. At least 1000 cells from three different experiments were monitored per experimental condition, except for diamide (20 cells), which has been characterized previously and is shown as reference. Results are shown as average values ± standard error or mean. # p< 0.005 when comparing Control vs H_2_O_2_ of C328H. (B) SW13/cl.2 cells stably transfected with RFP//vimentin wt or C328H were treated with the indicated agents, as above, and the morphology of the vimentin network was assessed by immunofluorescence and confocal microscopy. Images shown are overall projections. Cell contours are outlined in some images to illustrate vimentin network retraction from the cell periphery. Bars, 20 µm.

The insensitivity of vimentin C328H filaments to disruption by electrophiles was also confirmed in cells expressing the untagged proteins (Fig. 4B). Transfection of SW13/cl.2 cells with the RFP//vimentin C328H construct expressing untagged vimentin C328H and RFP as separate proteins, led to the formation of an extended filament network, similar to that of RFP//vimentin wt, as detected by immunofluorescence (Fig. 4B). Treatment of cells with various electrophilic agents, namely, diamide, HNE and DNI, elicited marked and differential alterations of the vimentin wt network. In particular, diamide induced a typical fragmentation of filaments into dots in virtually all cells expressing RFP//vimentin wt [18]. In turn, HNE typically provoked filament condensation near the center of the cell [58], and DNI elicited mixed alterations consisting in filament retraction from the cell periphery, disposition of filaments into parallel arrays and/or formation of thick filament bundles, alterations which affected approximately 75% and 68% of the cell population, respectively. In sharp contrast, vimentin C328H was virtually resistant to the filament fragmentation induced by diamide and clearly less susceptible to the alterations induced by HNE and DNI, since less than 26% of the cells displayed disrupted vimentin C328H upon treatment with these agents.

Taken together, these results show that vimentin C328H is markedly resistant to remodeling by diverse electrophiles and strengthen the role of C328 as a target for the action of various compounds disrupting vimentin organization.

### Expression of GFP-vimentin C328H blunts actin reorganization in response to electrophiles

As illustrated above, the vimentin network undergoes drastic rearrangements in response to oxidative and electrophilic stress. However, the significance of vimentin reorganization, i.e., whether it plays a protective, a damage-amplifying or a signaling role in cells, is not well understood. The ability of GFP-vimentin C328H to form short filamentous structures and its resistance to disruption by electrophiles provides a suitable framework to evaluate the impact of cysteine-dependent vimentin remodeling on other cellular events. Among them, actin organization is highly sensitive to electrophilic and oxidative stress (reviewed in [59–61]). Therefore, we monitored changes in f-actin as readout of the effects of electrophiles in cells expressing vimentin wt or C328H.

SW13/cl.2 cells expressing GFP-vimentin wt present a typical distribution of f-actin in cortical structures, focal adhesions and short cytoplasmic fibers of variable orientation (Fig. 5). Treatment of these cells with DNI elicited the formation of robust actin stress fibers.

**Figure 5.**
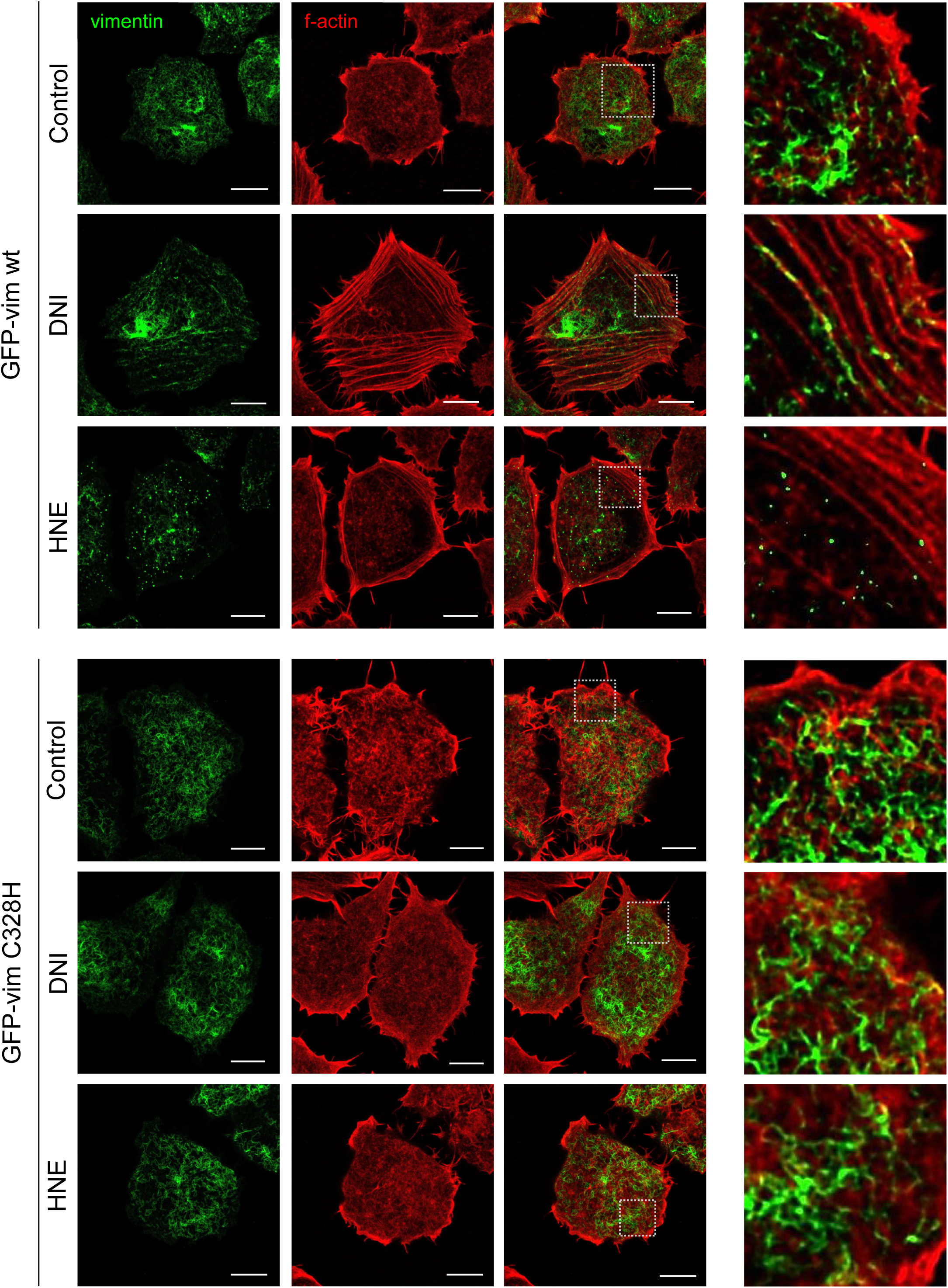
Effect of electrophiles on actin remodeling in cells expressing GFP-vimentin wt or C328H. SW13/cl.2 cells stably transfected with GFP-vimentin wt or C328H were treated with DNI or HNE, as above. After treatment, cells were fixed, f-actin was stained with phalloidin-TRITC and the distribution of GFP-vimentin (green) and f-actin (red) was assessed by confocal microscopy with the Lightning module. Images shown are single sections taken from the basal third of total cell height. Areas delimited by dotted squares in the merged images are enlarged at the right. Bars, 10 µm. Images shown are representative from at least three experiments for HNE and six for DNI.

Interestingly, f-actin staining revealed that the typical parallel arrays of GFP-vimentin wt short filaments induced by DNI treatment were actually aligned along actin fibers, as illustrated in the enlarged views of selected regions of the merged images (Fig. 5, right images). HNE treatment also elicited the appearance stress fibers, although in this case they were mainly subcortical and did not show any particular association with the disrupted vimentin structures.

Strikingly, DNI-elicited actin reorganization was markedly hampered in cells expressing GFP-vimentin C328H, which showed a drastic attenuation of the formation of stress fibers with respect to cells expressing GFP-vimentin wt (Fig. 5). Moreover, expression of GFP-vimentin C328H also impaired the formation of subcortical stress fibers in response to HNE.

The differential organization of actin in GFP-vimentin wt and C328H-expressing cells upon treatment with electrophiles could be confirmed by several types of analysis (Fig. 6). F-actin fluorescence intensity profiles along the lines drawn in images shown in Fig. 6A illustrate the appearance of a distinct saw-teeth pattern, typical of stress fibers, in GFP-vimentin wt-expressing cells treated with DNI, and the increases in subcortical fibers induced by HNE (Fig. 6A, arrowheads). Both features were absent in GFP-vimentin C328H-expressing cells, which presented an irregular f-actin profile without defined stress fibers similar to that of control cells. Moreover, the proportion of cells showing stress fibers in response to electrophile treatment was significantly higher in GFP-vimentin wt than in GFP-vimentin C328H expressing cells (Fig. 6B). Consistently, calculation of the variation coefficient of f-actin fluorescence intensity as an index of f-actin condensation into robust structures clearly showed that electrophiles elicited a significant increase of this parameter in cells expressing GFP-vimentin wt, which was abolished in GFP-vimentin C328H expressing cells (Fig. 6C). Finally, the degree of parallel alignment of stress fibers was estimated by calculating f-actin anisotropy [37]. Both DNI and HNE elicited significant increases (over 5-fold) of f-actin anisotropy in the presence of GFP-vimentin wt, which were blunted in GFP-vimentin C328H expressing cells (Fig. 6D). Notably, DNI also elicited stress fiber formation in MCF7 cells expressing GFP-vimentin wt, but this effect was abolished in cells transfected with GFP-vimentin C328H (Suppl. Fig. 3). Thus, electrophile-elicited actomyosin contractility appears to be precluded in cells expressing the electrophile-resistant vimentin C328H mutant.

**Figure 6.**
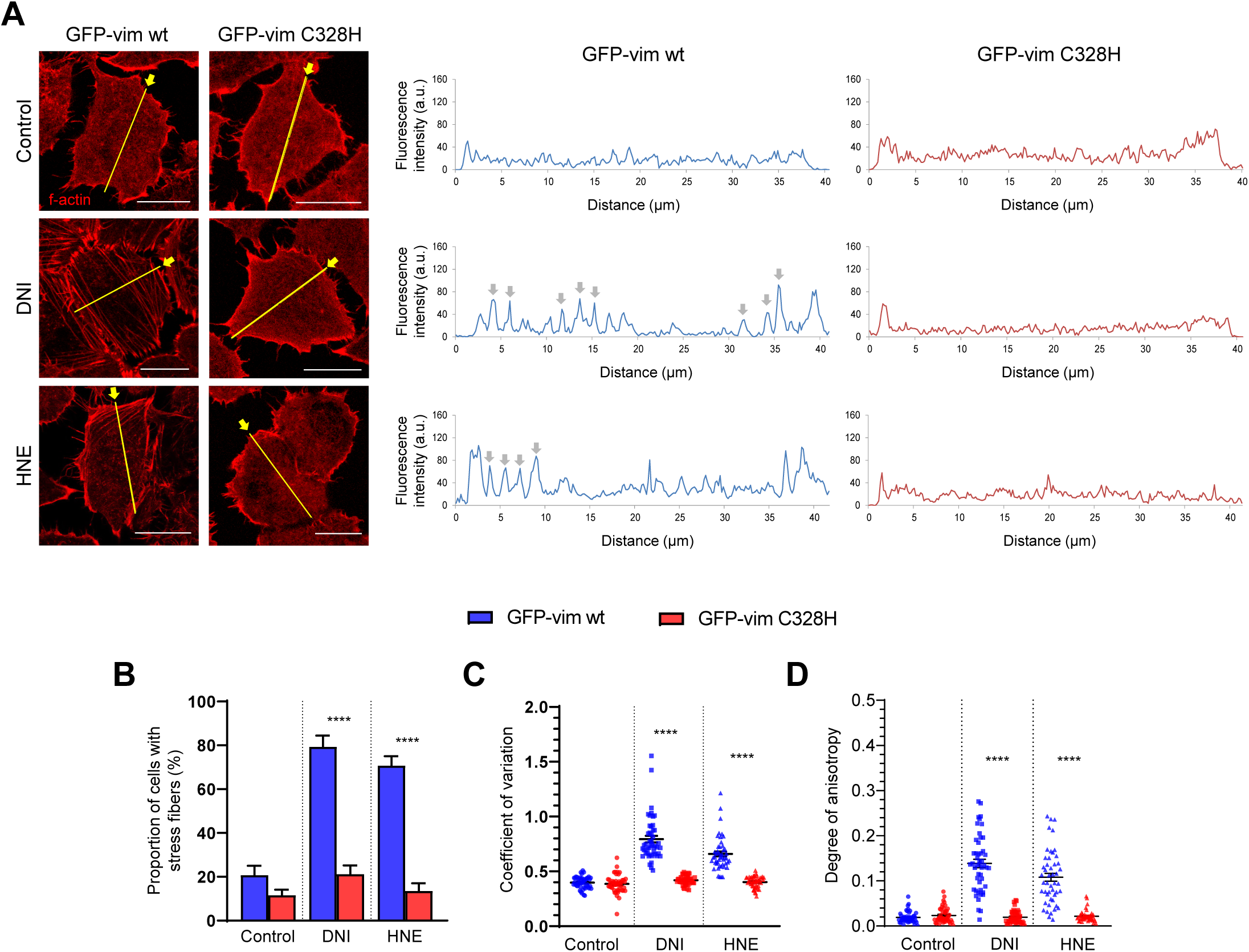
Analysis of the distribution of f-actin in cells expressing GFP-vimentin wt or C328H. SW13/cl.2 cells were treated with DNI or HNE as in Fig. 5, and f-actin was stained with phalloidin-TRITC. (A) Fluorescence intensity profiles of f-actin distribution were obtained along the lines drawn on images in the direction indicated by arrows. Profiles shown are representative from those obtained from at least 45 cells per experimental condition from three independent experiments. (B) The proportion of cells displaying stress fibers was assessed by visual inspection by monitoring a total of 450 cells per experimental condition from three different experiments. (C) The coefficient of variation of the f-actin signal was calculated from the values of the fluorescence intensity plots obtained in (A). (D) The degree of anisotropy of f-actin structures was quantitated from three independent experiments, totaling at least 48 cells per experimental condition. Results are shown as average values ± S.E.M. Bars, 20 μm.

These results suggest that the presence of a cysteine residue at position 328 in vimentin is important for cytoskeletal reorganization upon electrophilic stress. This could be due to the particular susceptibility of C328 to modification by electrophiles, either directly (adduct formation) and/or indirectly (oxidative modifications). Indeed, HNE is well known to modify C328 in several experimental models related to pathophysiological conditions [35, 62, 63]. However, to the best of our knowledge, modification of proteins in intact cells by DNI has not been reported yet. To tackle this point, here we have assessed free cysteine availability in vimentin in control and DNI-treated cells by using biotinylated iodoacetamide and biotinylated maleimide labeling (Suppl. Fig. 4). We have observed a decrease in this parameter in DNI-treated cells, which is only partially reversed by incubation of cell lysates with DTT (Suppl Fig. 4), indicating that DNI elicits both reversible and irreversible modifications of vimentin C328.

### A GFP-vimentin C328D mutant neither elicits nor blocks actin remodeling in response to DNI

Although the results shown above support the occurrence of vimentin cysteine modifications, it is not clear how these modifications can impact actin remodeling. Two main possibilities can be considered. 1) Vimentin perturbations at C328 could affect actin remodeling by actively contributing to actin stress fiber formation (positive effect). 2) Alternatively, C328 modification could impair a repressive function of vimentin on acto-myosin contractility, therefore allowing stress fibers to form (neutralization of a negative effect).

Cysteine to aspartate mutants have been previously used as mimics of certain oxidized forms of cysteine residues [64, 65]. Therefore, to test the first possibility, we explored the distribution of a GFP-vimentin C328D mutant and its impact on actin organization under control conditions and upon electrophilic stress (Fig. 7). Interestingly, GFP-vimentin C328D was not able to form regular squiggles or short filaments in cells. Instead, bright cytoplasmic accumulations were observed (Fig. 7), indicating that an aspartate residue at this position is not tolerated for assembly of the GFP-vimentin construct. Notably, expression of GFP-vimentin C328D was not associated with an increase in the proportion of cells displaying stress fibers under control conditions, which represented 20-25% of the cell population, a proportion similar to that present in cells expressing GFP-vimentin wt or in non-transfected cells (Fig. 6 and 7).

**Figure 7.**
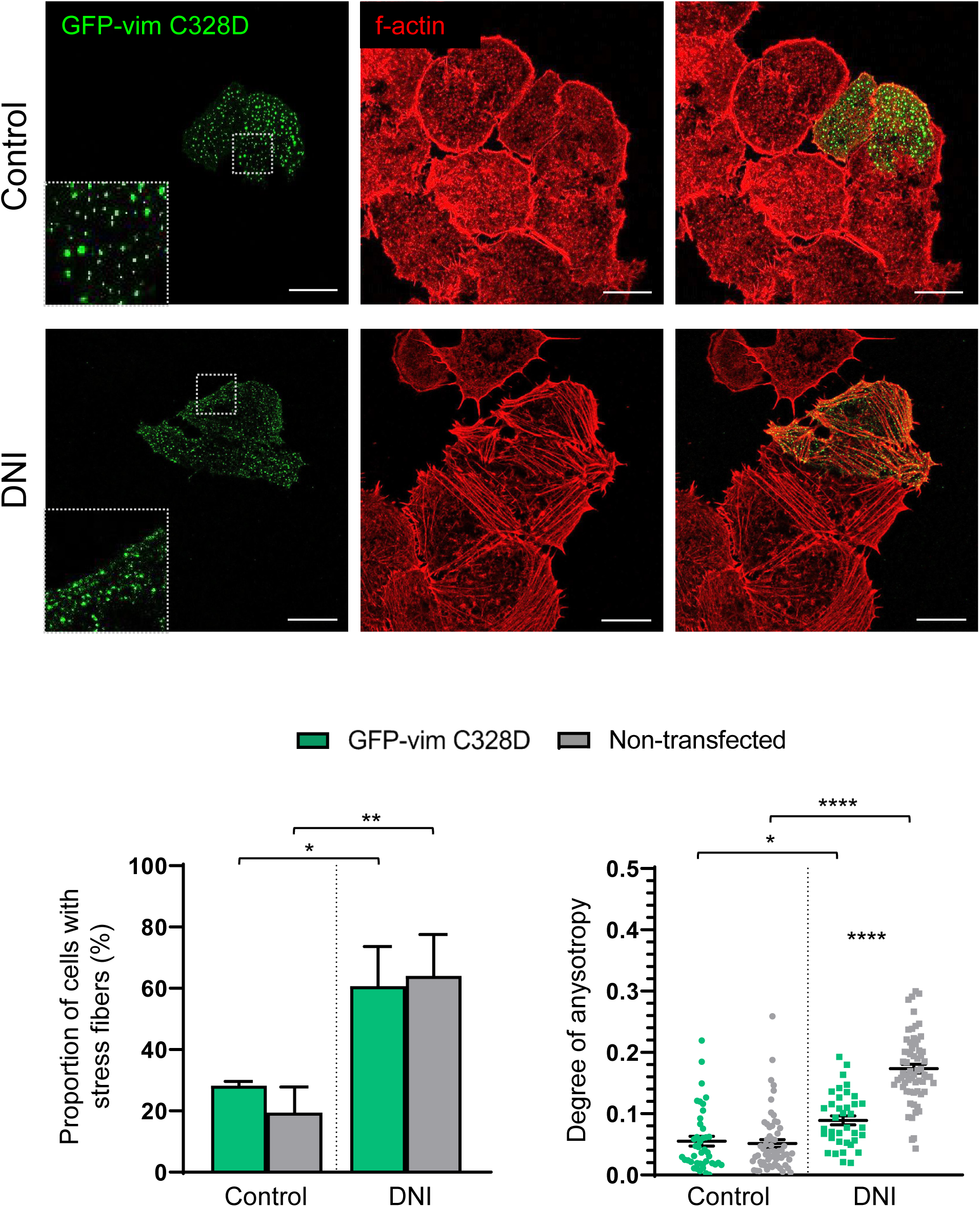
Effect of DNI on actin remodeling in cells expressing a GFP-vimentin C328D mutant, in comparison with non-transfected cells. SW13/cl.2 cells were transiently transfected with GFP-vimentin C328D and treated with DNI, as indicated. Cells were fixed and f-actin was stained with phalloidin-TRITC. The distribution of GFP-vimentin C328D (green) and actin (red), in cells expressing the GFP-vimentin C328D protein and in non-transfected cells from the same experiments, was monitored by confocal microscopy. Single sections were taken from the lower third of cells and individual channels and merged images are shown. The proportion of cells with stress fibers and the degree of anisotropy of f-actin structures were obtained as above from at least from three different experiments totaling at least 40 cells per experimental condition, and are depicted in the lower graphs. Results are shown as average values ± S.E.M. Bars, 20 µm.

Therefore, this “pseudo oxidation mutant” did not trigger actin remodeling *per se*, suggesting that at least some oxidative modifications of C328 may not be sufficient to elicit actin reorganization. Treatment with DNI did not alter the punctate pattern of GFP-vimentin C328D, although vimentin dots adopted a more linear distribution. Interestingly, staining of f-actin revealed that GFP-vimentin C328D did not block the induction of robust stress fibers by DNI treatment, when compared to non-transfected cells (see for instance image at bottom right in Fig. 7). In fact, numerous GFP-vimentin C328D dots were aligned along the trajectories of stress fibers. Quantitation of these effects showed that the proportion of cells displaying stress fibers in response to DNI was similar in cells expressing GFP-vimentin C328D and in non-transfected cells. Moreover, DNI-elicited stress fiber anisotropy was slightly but consistently higher in non-transfected cells than in cells expressing GFP-vimentin C328D. Taken together, these observations suggest that C328 modification does not imply a “gain of function” on actin remodeling.

### The presence of vimentin negatively modulates serum-elicited stress fibers

To explore whether vimentin exerts a negative effect on acto-myosin contractility, we evaluated its impact on serum-elicited stress fibers. Consistent with the observations in non-transfected cells shown in Fig. 7, only a small proportion of vimentin-deficient SW13/cl.2 cells displayed stress fibers under normal cell culture conditions or after serum starvation for 24 h (Fig 8). Acute stimulation of starved cells with serum elicited robust stress fibers. Importantly, this induction was markedly attenuated in cells expressing vimentin wt, as denoted by a significant decrease both in the proportion of cells displaying stress fibers and in the anisotropy of the f-actin signal (Fig. 8, graphs). Of note, serum deprivation or addition did not appreciably alter vimentin organization (Fig. 8, insets). Taken together, these results point towards the second possibility raised above, that is, the existence of a suppressive effect of vimentin on actin remodeling that could be counteracted by C328 perturbation by electrophiles.

**Figure 8.**
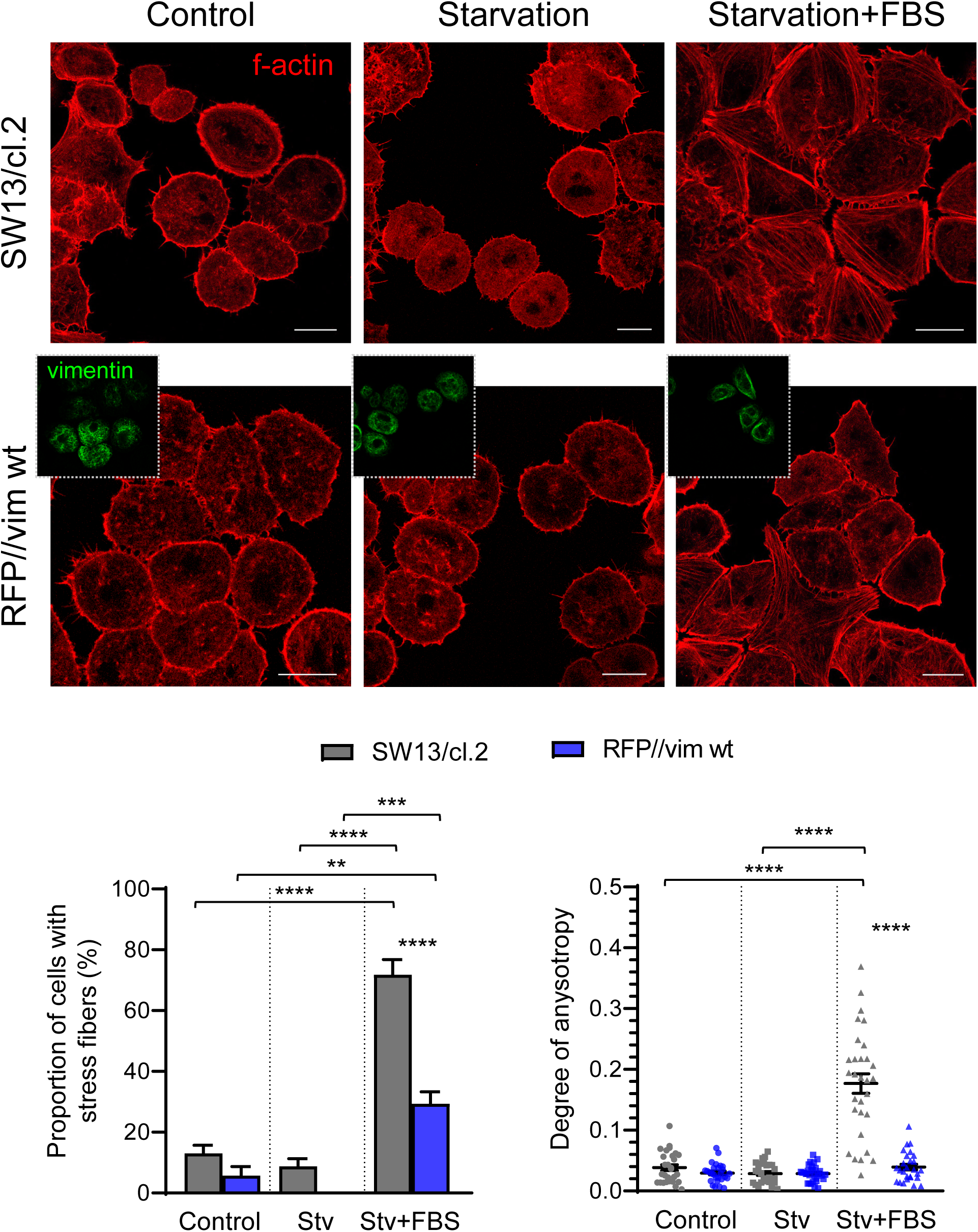
Effect of vimentin on the formation of actin stress fibers in response to serum stimulation. SW13/cl.2 cells, non-transfected (upper row) or stably transfected with vimentin wt (RFP//vim wt, lower row) were kept under control culture conditions (left images) or starved for 24 h (middle images). Serum-starved cells were stimulated with 10% (v/v) FBS for 30 min (right images). After treatments, cells were fixed and processed for vimentin and f-actin detection by immunofluorescence with the V9 antibody and Phalloidin-TRITC staining. Insets in the lower row display vimentin staining. The bottom-right image clearly shows the attenuation of stress fibers formation in vimentin-positive-cells in comparison with nearby cells not expressing vimentin. The graphs depict the proportion of cells displaying stress fibers (left), and the degree of anisotropy of the f-actin signal (right). The bar corresponding to the proportion of RFP//vim cells with stress fibers after starvation is not visible because the value is zero. Results from at least 30 determinations are shown as average values ± S.E.M Bars, 20 µm. Stv, starvation.

### Involvement of the RhoK pathway in DNI-induction of stress fibers and its modulation by vimentin

A classical/well characterized pathway for stress fiber induction involves the activation of RhoA GTPase and its downstream effector RhoK, which in turn regulates actin organization and actomyosin contractility through direct and indirect mechanisms, including an increase in myosin phosphorylation and f-actin stabilization (reviewed in [66, 67]). Importantly, vimentin has been reported to exert a negative effect on stress fibers under control conditions in U2OS cells by attenuating the RhoA pathway [68]. This prompted us to assess the putative involvement of the RhoA pathway in the effects of DNI on actin organization (Fig. 9). Interestingly, pretreatment with the RhoK inhibitor, Y27632, abolished DNI induction of stress fibers (Fig. 9A), thus pointing to the Rho pathway as the main mechanism for this effect.

**Figure 9.**
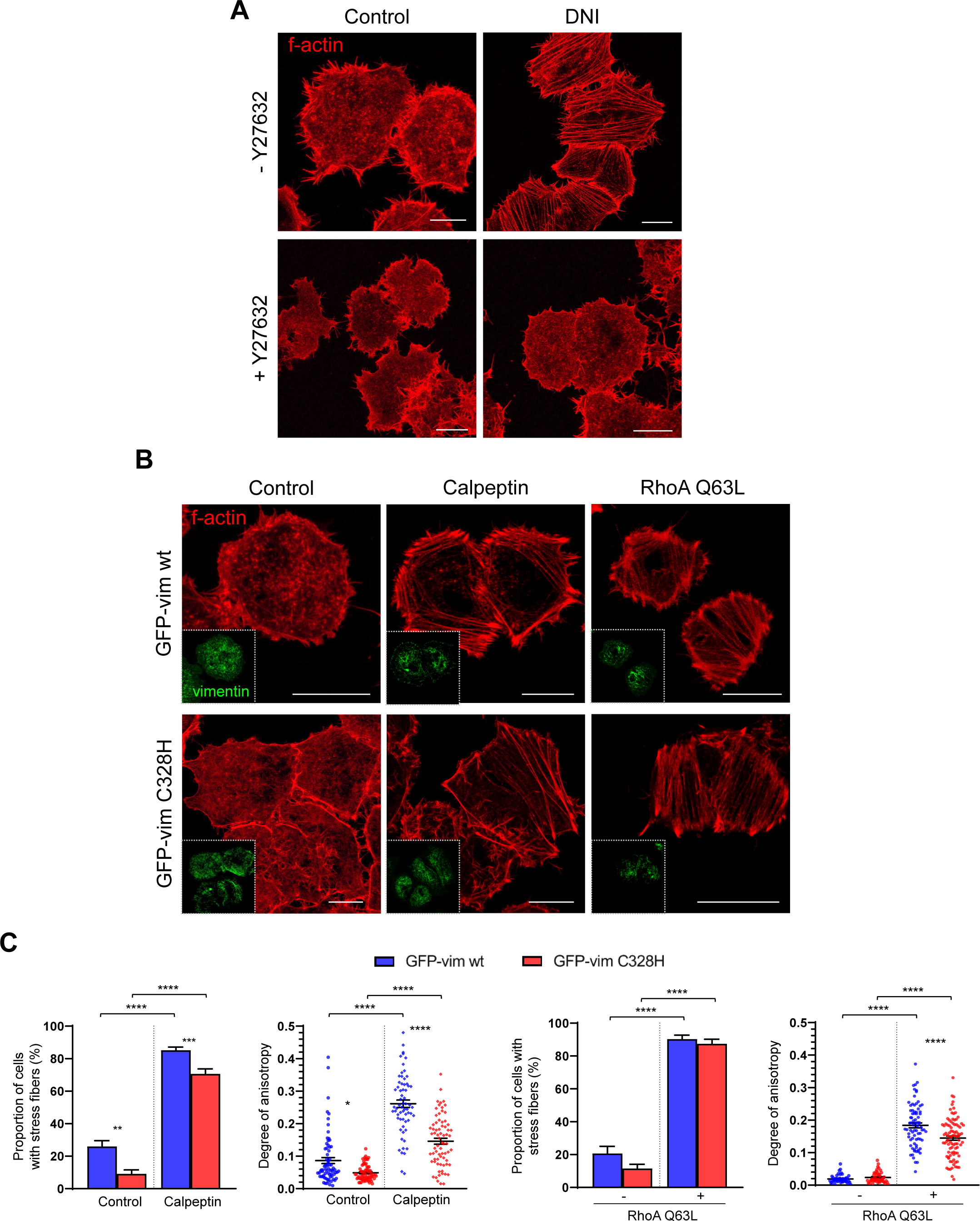
Effect of the modulation of the RhoA-RhoK pathway on DNI-elicited actin remodeling. (A) SW13/cl.2 cells were incubated in the absence or presence of Y27632, and subsequently treated with DNI, as indicated in the experimental section. (B) Cells expressing GFP-vimentin wt of C328H were treated with calpeptin or transiently transfected with RhoA Q63L, as previously described. Panels (A) An d (B) show cells were fixed and stained with phalloidin-TRITC and visualized by confocal microscopy. Insets show the presence of the corresponding vimentin construct in the cells monitored. Images shown are overall projections and are representative from three experiments with similar results, and quantitation of these effects is shown in (C). The minimum number of cells monitored per experimental condition was 150 for calculation of the proportion of cells displaying stress fibers and 60 cells to obtain anisotropy values. Bars, 20 µm.

Next, we explored whether activation of RhoA by alternative means, overcame the negative effect of vimentin C328H on stress fiber formation. The compound calpeptin is a phosphatase inhibitor reported to act as a RhoA activator independently from the generation of ROS [69]. We observed that calpeptin potently induced stress fibers, both in GFP-vimentin wt and in GFP-vimentin C328H expressing cells, as indicated by the proportion of cells displaying stress fibers and their degree of anisotropy. Nevertheless, the extent of induction was more moderate in cells expressing GFP-vimentin C328H (quantitated in Fig. 9C). Moreover, transfection with a constitutively active RhoA construct, namely, RhoA Q63L, elicited a massive formation of robust stress fibers, frequently spanning all the length of the cell (Fig. 9B), which occurred in a similar proportion of cells expressing GFP-vimentin wt or C328H (Fig. 9C).

Constitutively active RhoA also elicited an increase in f-actin anisotropy, although less marked in GFP-vimentin C328H-expressing cells. Therefore, GFP-vimentin C328H was not able to block stress fiber formation induced by active RhoA (Fig. 9B and C). Taken together, these results indicate that vimentin C328H is particularly effective at blocking stress fiber formation elicited by certain electrophiles, and this inhibition seems to take place upstream of RhoA activation (see Fig. 11).

### Inhibition of DNI-elicited stress fibers requires electrophile-resistant vimentin filamentous structures

The existence of vimentin as filamentous structures is important for several functions of the protein, including its role in metastasis [70], interplay with RhoA [68], and the upregulation of transcription factor TWIST1, which is related to epithelial-mesenchymal transition [71]. The results shown above indicate that both, the elongation-defective vimentin construct GFP-vimentin C328D and the electrophile-susceptible GFP-vimentin wt, do not block the formation of stress fibers in response to DNI, whereas electrophile-resistant vimentin C328H filamentous structures blunt actin reorganization. To confirm that filamentous, undisrupted vimentin is necessary to hamper DNI-elicited stress fiber induction, we employed additional constructs. The vimentin C328A mutant provides an interesting case since the GFP-fusion version is unable to elongate and forms only dots in vimentin-deficient cells (see Fig. 3), whereas its untagged version assembles into extended filaments that are resistant to disruption by the electrophile diamide [16]. As shown in Fig. 10A, as well as earlier in Fig. 4, electrophile-susceptible untagged vimentin wt reorganized into parallel arrays upon treatment with DNI, which turned out to be disposed along and interweaving with actin stress fibers (Fig. 10A). In contrast, filaments formed by electrophile-resistant untagged vimentin C328A were not altered by DNI and were able to counteract the induction of stress fibers, blunting the increase in the proportion of cells showing stress fibers and in f-actin anisotropy (Fig. 10B). Remarkably, GFP-vimentin C328A dots did not block DNI-elicited stress fibers, which were detected in nearly 75% of the cells expressing this construct, and displayed increased anisotropy with respect to the control (Fig. 10C). In turn, DNI did not alter the punctate pattern of GFP-vimentin C328A, though it provoked its distribution into linear alignments, as observed for GFP-vimentin C328D (Fig. 10C).

**Figure 10.**
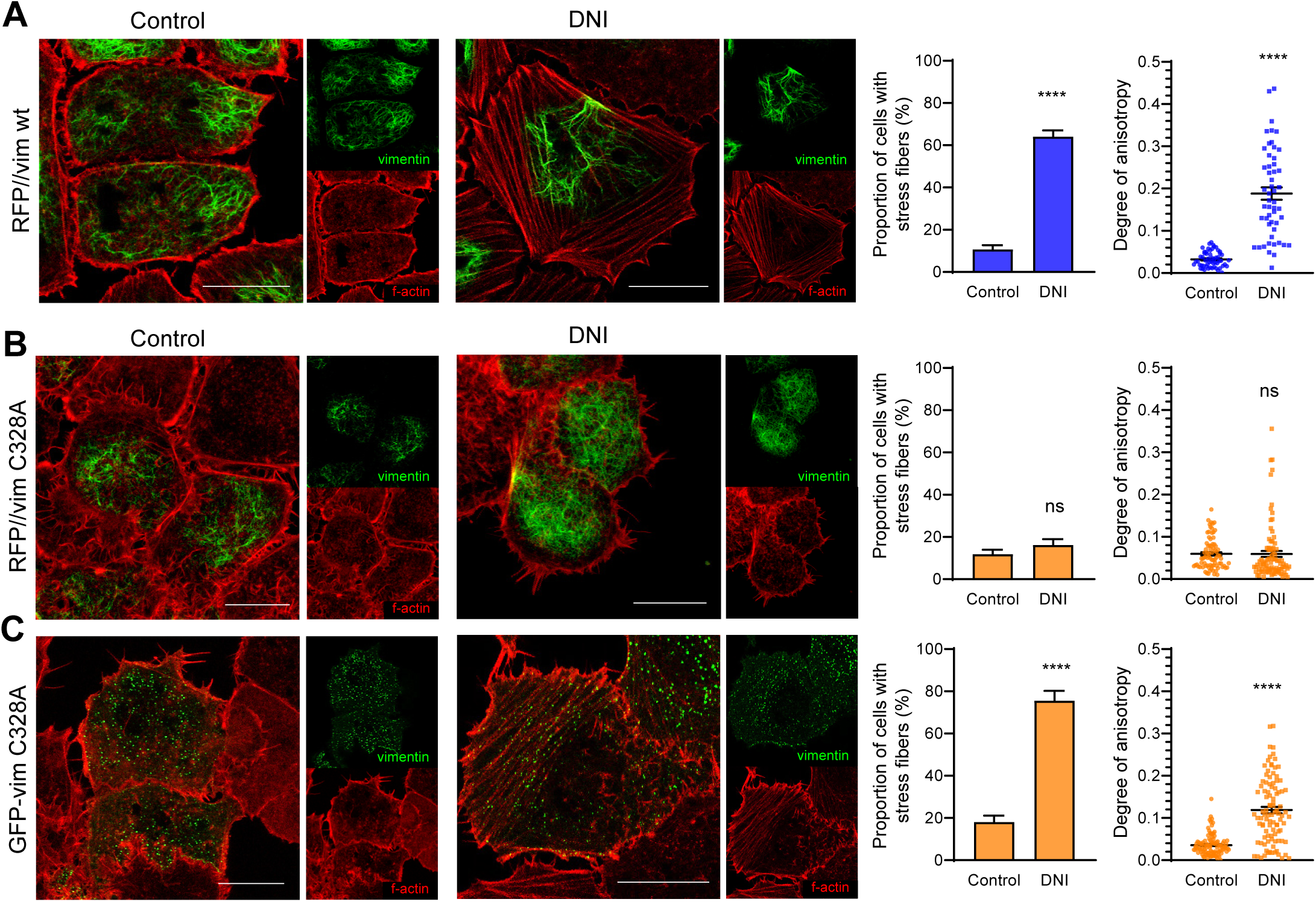
Role of vimentin filamentous structures in the modulation of actin remodeling in response to DNI. SW13/cl.2 cells stably expressing, (A) RFP//vimentin wt, (B) RFP//vimentin C328A or (C) GFP-vimentin C328A, were treated with DNI as indicated, fixed and stained with phalloidin-TRITC for monitoring actin distribution. In (A) and (B) cells vimentin was visualized by immunofluorescence, and in (C) by direct observation of GFP fluorescence. Merged images of single sections and individual channels (small panels at the right) are shown. Bars 20 µm. Graphs on the right depict the proportion of cells displaying stress fibers, and the degree of anisotropy of the f-actin signal for every experimental condition. A minimum of 170 and 50 cells per experimental condition were monitored for calculation of the proportion of cells displaying stress fibers and anisotropy values, respectively. Results are average values ± S.E.M.

Taken together, the results shown above suggest that the ability of vimentin to block DNI-elicited actin stress fibers requires its presence in filamentous structures resistant to disruption by this electrophile. In this scenario, cysteine-mediated vimentin disruption by electrophiles could be a pre-requisite for stress fiber formation. A summary of these findings, presented in Fig. 11, supports the existence of a vimentin-actin interplay under stress which is dependent on vimentin network remodeling mediated by modification of C328.

**Figure 11.**
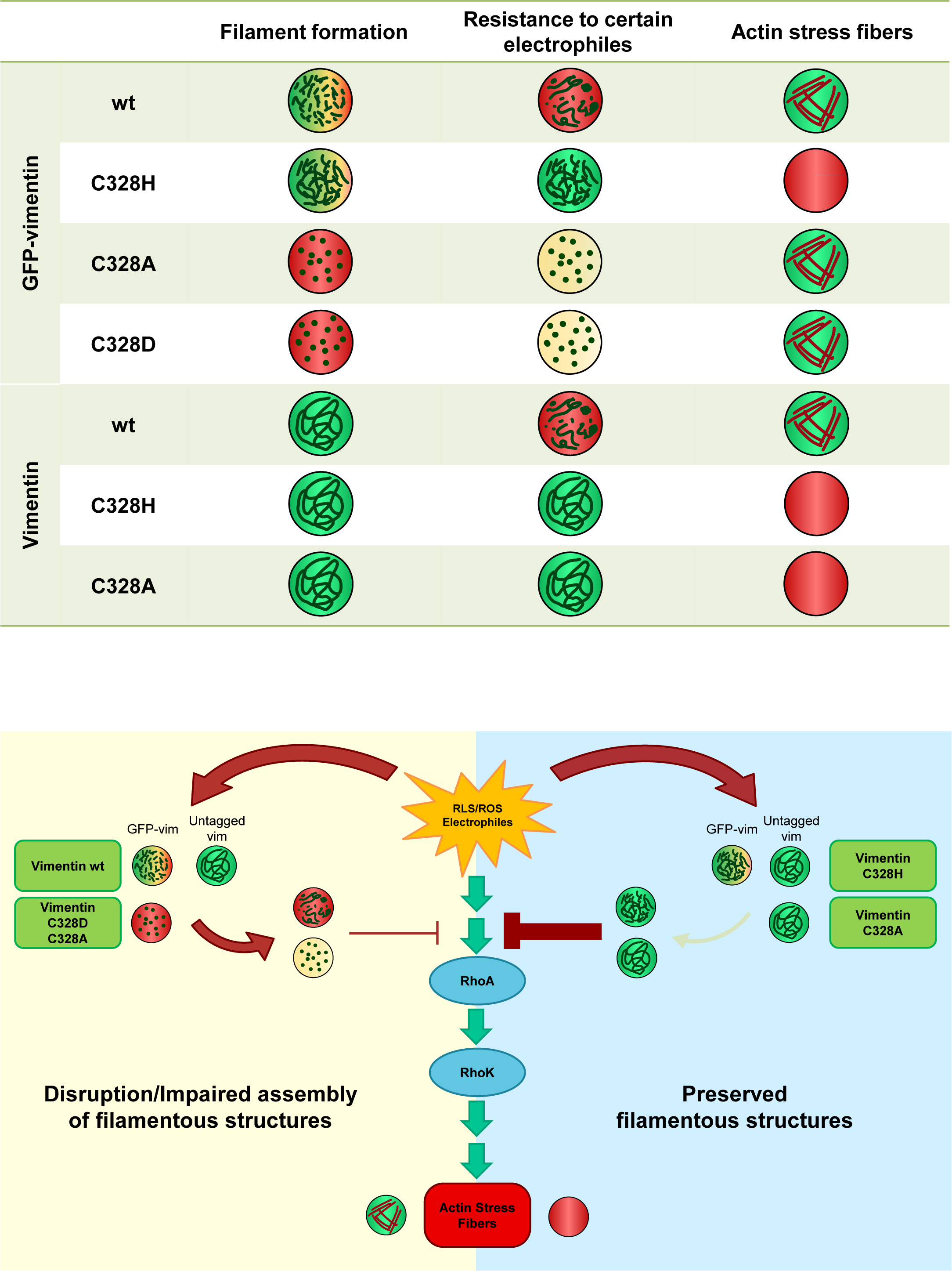
Summary of the behavior of the various vimentin forms used in this study and their potential involvement in the regulation of actin stress fibers upon treatment with certain electrophiles. (A) The behavior of vimentin GFP fusion constructs (GFP-vimentin) and untagged full-length vimentin wt or mutants (vimentin) is schematized. Circles with a green background indicate preserved function, whereas a red background indicates impairment of the corresponding process. Yellow circles are used in the case of vimentin constructs with defective assembly and/or elongation. Mixed patterns are indicated by a combination of the corresponding color codes. (B) Hypothetical model for the involvement of vimentin in the regulation of actin stress fibers elicited by certain electrophiles. Electrophiles such as DNI elicit actin stress fibers through the activation of the Rho/RhoK pathway (center), and cause the disruption of vimentin wt (left), but not of certain vimentin mutants (right). Undisrupted, electrophile-resistant filamentous vimentin structures would block RhoA/RhoK activation, thus attenuating the induction of stress fibers by electrophiles (right). Thin arrows or connectors indicate steps/processes that are attenuated under the experimental conditions used. RLS/ROS, reactive lipid or oxygen species.

## 4. Discussion

Cysteine residues are key players in redox and electrophile signaling, due to their particular reactivity and their capacity to undergo modifications incorporating widely diverse moieties, which may differentially affect the structure and function, and even the subcellular localization of the target proteins (reviewed in [40]). Cytoskeletal proteins, including actin, tubulin and vimentin, have been frequently identified as targets for cysteine modifications, which in some cases have been found to be of regulatory importance [34, 72–74]. Vimentin organization is highly responsive to modulation by oxidants and electrophiles, and the single cysteine residue within its sequence, C328, has been shown to undergo various modifications and to play a key role in vimentin reorganization in response to some of these agents, both *in vitro* and in cells [16, 18, 24, 29, 31, 32, 34, 35]. Therefore, a role for this cysteine residue in redox and stress sensing has been put forward [12, 16]. Nevertheless, to date, the downstream consequences of C328-mediated vimentin remodeling on cell behavior have not been elucidated. Here, we show that perturbations at the 328 position lead to structure-dependent vimentin remodeling, suggesting that different modifications may elicit distinct consequences. Moreover, our results unveil that C328-dependent vimentin susceptibility to disruption is necessary for full actin reorganization in response to various electrophiles. Therefore, C328 emerges as a gatekeeper and a tunable hinge for thiol/redox-mediated modulation of vimentin, which appears as a novel modulatory element in the pathway transducing signals from reactive species into actin responses.

In early works we identified C328 of vimentin as a selective site for modification by cyclopentenone prostaglandins [34, 75], and later noted that the presence of this residue was required for vimentin remodeling in response to different reactive species, endogenous mediators or exogenous electrophiles [16]. Interestingly, accumulating evidence from *in vitro* studies indicates that purified vimentin bearing different modifications at C328 engages in morphologically distinct assemblies. Indeed, disulfide crosslinking affecting approximately 50% of the protein impairs filament formation leading to aggregates [18, 29], whereas quantitative glutathionylation not only impairs filament elongation but severs preformed filaments [31]. In contrast, S-nitrosation is associated with minor effects on assembly [31], and mild oxidation or lipoxidation increase filament width [18]. Therefore, structurally different modifications at C328 appear to result in diverse functional outcomes *in vitro*.

However, this picture becomes more complex in the cellular context, where an array of modifications can coexist or evolve in a time-dependent fashion, leading to a wide diversity of vimentin proteoforms, even after exposure to a single agent. For instance, as we reported previously in cardiac cells, exposure to the peroxynitrite donor SIN-1 elicits multiple PTMs on vimentin in a dynamic fashion [24]. Indeed, many electrophilic compounds can induce oxidative stress or stimulate the generation of reactive species, potentially leading to secondary modifications, such as cysteine oxidations or nitrosation [24]. Additionally, other cellular factors, including metals, small molecules and antioxidants can influence cysteine reactivity [16].

Moreover, interplay between direct vimentin modification by oxidants and electrophiles and other redox-regulated modifications, such as phosphorylation or acetylation, also occurs (reviewed in [12, 19]), increasing this complexity.

The observations gathered herein show that, indeed, the cellular vimentin network undergoes distinctive remodeling in response to different agents, and in some cases, adopts peculiar patterns, as in the case of cyclohexenone-induced curly filaments or of DNI-elicited parallel linear arrays. However, these complex rearrangements are likely the result of multiple PTMs affecting not only vimentin but also other cellular targets acting on the vimentin network. In fact, as stated above, most electrophilic compounds can modify several cellular targets and/or elicit the generation of reactive species, contributing to PTM crosstalk [53, 54, 55].

Indeed, we have obtained evidence of an increased formation of protein carbonyls in cells treated with HNE, cyclohexenone and DNI (Suppl. Fig. 1), as well as indications for reversible and irreversible modification of C328 by the latter (Suppl. Fig. 4). These factors make it virtually unfeasible to ascribe specific structural perturbations to functional outcomes in cells, and hence to focus on effects from a single vimentin proteoform.

To circumvent this complexity, we studied the organization of vimentin cysteine mutants bearing amino acid substitutions that may be structurally reminiscent of some of the modifications elicited by the electrophilic compounds used. Indeed, introduction of mutations that mimic or preclude certain PTMs is a strategy widely used to perturb only the protein of interest, in our case, vimentin. Our results clearly show that, in cells, mutations at the 328 position differentially impact the morphology of the vimentin network depending on the new side chain introduced, indicating that alterations at the cysteine site may be sufficient to elicit distinct vimentin reorganizations. Remarkably, although most of the mutants generated markedly altered vimentin structures, the GFP-vimentin C328H mutant formed nearly normal, but somehow more robust, squiggles than GFP-vimentin wt in SW13/cl.2 cells, and very robust filaments and bundles in MCF7 cells. Moreover, untagged vimentin C328H formed an extended filament network, similar to wt. For these reasons, we studied the C328H mutant in more detail.

Interestingly, when assessing the response to oxidants and electrophiles, we observed a nearly complete resistance of vimentin C328H to disruption by these agents. This protection is particularly striking in the cases of diamide and DNI, which elicit drastic rearrangements of both GFP-vimentin wt and untagged vimentin wt structures, whereas vimentin C328H squiggles and filaments maintain their regular morphology. Vimentin C328H resistance to these agents could arise from the lower nucleophilicity of histidine, compared to the cysteine residue, and therefore its lower reactivity towards electrophiles [76]. Histidine residues can also undergo oxidation, although the reaction constants with certain oxidants are higher than those of cysteine residues [77], and they cannot undergo the straight-forward oxidation characteristic of the sulfur center [78]. The slightly lower resistance of C328H to disruption by H_2_O_2_, compared to other treatments, could be due to the ability of this agent to oxidize histidine residues [79] or to generate other reactive species, such as the hydroxyl radical, which can induce lipid peroxidation, giving rise to various electrophilic lipids, and subsequent protein lipoxidation [80, 81]. Nevertheless, our results clearly show that, in spite of the broad reactivity of the compounds used and their ability to modify multiple cellular targets and protein residues, the presence of a cysteine residue at the 328 position, and therefore the high susceptibility of the thiol group to direct or indirect modification, is critical for vimentin remodeling to occur.

Moreover, considering the differential organization of the various cysteine mutants, it could be proposed that C328 could act as a “plug” accepting diverse moieties, which would result in distinct functional outcomes [12, 82]. The mechanisms linking perturbations at the cysteine site with particular vimentin rearrangements could be related to the location of C328 at a position critical for filament assembly and/or elongation, perhaps close to the site of imbrication of adjacent vimentin subunits (or unit length filaments, ULF), in such a way that bulky modifications would have a marked impact on the morphology of the network, as previously discussed [12]. Indeed, C328 glutathionylation of purified vimentin precludes longitudinal assembly of vimentin ULF, thus impairing elongation, as evidenced by highly sensitive hydrogen-deuterium exchange mass spectrometry experiments [31].

Although modification of C328 by electrophiles appears critical for vimentin remodeling, it is not known whether it impacts other cellular responses. The C328H mutant provides a means of exploring how electrophiles affect cellular behavior in the absence of vimentin reorganization, therefore, shedding light on the importance of vimentin disruption for other cellular effects of electrophiles. Several processes known to be affected by vimentin could be monitored, including organelle positioning, cell adhesion and migration, or cytoskeletal crosstalk. Of these, we focused on actin remodeling, given the close interplay between vimentin and actin. Previous studies demonstrated that actin structures modulate the traffic of vimentin precursors and filaments, whereas vimentin modulates the organization of actin in various structures, including actomyosin arcs and stress fibers, cortical actin and focal adhesions [8, 68, 83]. Notably, the actin cytoskeleton itself is susceptible to regulation by oxidants and reactive agents [72], both through direct and indirect mechanisms. In particular, H_2_O_2_ can induce the formation of actin stress fibers in a cell type-dependent manner [84, 85], whereas reactive lipids can elicit, disrupt or remodel stress fibers [37, 80]. Surprisingly, we found that the potent actin reorganization elicited by the electrophiles DNI and HNE in cells expressing vimentin wt, was virtually blunted in cells bearing the electrophile resistant vimentin C328H. Therefore, just this point mutation in vimentin drastically altered actin responses.

Several possibilities can be envisaged that could contribute to this effect. First, electrophiles could elicit a particular modification of vimentin C328 with a positive role in stress fiber formation. This possibility cannot be ruled out, although vimentin C328D, mimicking cysteine oxidation, does not induce stress fibers *per se*. On the other hand, vimentin wt could exert an indirect effect by scavenging certain reactive species or mediators with negative impact on stress fiber formation. Additionally, vimentin itself could play a negative role on stress fibers, which would be neutralized by electrophile-elicited vimentin disruption. Indeed, here we have shown that the presence of vimentin dampens the induction of stress fibers by serum stimulation. These results are in agreement with previous reports of a higher abundance of stress fibers in vimentin and plectin knockout cells than in their wt counterparts, which also point to the inhibition of actin organization by vimentin [68, 86]. Therefore, although other interpretations are possible, our results are consistent with vimentin exerting a negative effect on stress fiber formation, which would be relieved by its disruption by electrophiles, allowing full actin remodeling. Such a disruption would occur in cells expressing vimentin wt, in which the presence of C328 makes the protein highly susceptible to modification. Conversely, this sequence of events would be impaired in cells expressing the less nucleophilic vimentin C328H mutant, which is highly resistant to disruption by oxidants and electrophiles. This hypothesis is summarized in Fig. 11.

A major mechanism for stress fiber formation is the activation of the RhoA-RhoK pathway. Indeed, we have observed that treatment with a RhoK inhibitor blunts stress fiber formation in response to DNI, whereas a RhoA constitutively active mutant or treatment with calpeptin, potently elicit stress fiber formation in the presence of either GFP-vimentin wt or C328H. Therefore, the blocking effect of vimentin C328H appears to occur upstream of RhoA activation. The mechanism of vimentin inhibitory effect on RhoA activation could be multiple. In fact, a negative regulation of RhoA activity either through modulation of RhoA GEFs or of focal adhesions [68, 86] has been proposed. Nevertheless, it also needs to be taken into account that oxidants and electrophiles can regulate several elements of the RhoA/RhoK pathway, including RhoA itself, as well as other GTPases, which in turn could influence actin organization, and even vimentin phosphorylation and disposition [37, 84, 87–90].

Additional insight into the mechanism of vimentin-actin interplay has been previously obtained from the study of elongation-defective mutants, which showed that inhibition of actomyosin contractility required the presence of vimentin in filaments [68, 91]. Indeed, here we have observed that GFP-vimentin C328D and C328A, which are unable to form filamentous structures, do not preclude stress fibers induction by electrophiles. Conversely, untagged vimentin C328A, which forms long filaments resistant to disruption by cysteine-targeting agents, potently suppresses DNI-elicited stress fibers. In this scenario, under certain types of stress, the ability of vimentin wt filamentous structures to attenuate actomyosin contractility would be overcome by cysteine modification-mediated network disruption. This is clearly illustrated in cells expressing untagged vimentin wt, which forms long filaments that retract from the cell periphery or form accumulations upon treatment with DNI, allowing the formation of robust stress fibers, as depicted in Fig 10A. The behavior of the various vimentin constructs employed in our study, in terms of assembly, resistance to DNI-elicited disruption and ability to block DNI-induced actin stress fibers is summarized in Fig. 11A. Considering these observations, it could be hypothesized that induction of stress fibers by certain reactive compounds would be favored by cysteine modification and disruption of vimentin, potentially disabling its negative effect on the RhoA/RhoK pathway, therefore allowing activation (Fig. 11B). In contrast, disruption-resistant mutants, C328H or C328A, would persistently block this pathway. This encompasses the possibility that, under some circumstances, vimentin C328 could be acting as a redox or stress responsive element to influence RhoA signaling.

Nevertheless, other mechanisms and interactions need to be considered. For instance, vimentin wt and the various vimentin proteoforms bearing cysteine modifications or mutations could be differentially involved in protein-protein interactions with impact on the actin cytoskeleton; among them, those involving cytolinkers, filamin or plectin, which act as bridges between intermediate filaments and microfilaments or actin-regulating proteins [86, 92, 93].

Indeed, both plectin and filamin contain cysteine residues that have been identified as targets for nitrosation or other redox modifications [94, 95]. Therefore, cysteine-deficient or modified vimentin proteoforms could establish interactions with plectin or other cytolinkers, different from those established by vimentin wt, resulting in alteration of the cytoskeletal redox crosstalk.

In spite of the evident complexity of vimentin regulation and vimentin-actin interplay, taken together, our results show that vimentin single cysteine residue plays a regulatory role, acting as a hub for diverse modifications mediating the structure-dependent reorganization of vimentin in response to oxidants and electrophiles. Moreover, C328 modifications resulting in network disruption release a vimentin “break” on actin remodeling, allowing stress fiber formation in response to certain electrophiles. Therefore, these observations highlight C328 as a sensor, i.e., a residue that detects changes in the environment, such as the presence of certain reactive agents, and elicits or influences cellular responses downstream of vimentin, and renew the interest in finding C328-binding molecules as potential modulators of vimentin function.

## 5. Concluding remarks

The intermediate filament protein vimentin plays an important role in cytoskeletal crosstalk and is arising as a “privileged” target for oxidants and electrophiles. Our results show that these reactive compounds markedly reorganize vimentin wt. Remarkably, this remodeling is drastically attenuated in cells expressing cysteine-deficient vimentin mutants, even in the presence of extensive modification(s) of other cellular targets by these compounds. Moreover, perturbations introduced specifically at the 328 site through mutagenesis are sufficient to remodel the vimentin network. Regarding cytoskeletal crosstalk, C328-dependent vimentin remodeling appears to be required for the induction of actin stress fibers in response to certain electrophiles. Indeed, simply expressing electrophile-resistant vimentin cysteine mutants is sufficient to blunt stress fiber formation, even in the presence of electrophiles capable of eliciting multiple cellular effects. Therefore, the single vimentin cysteine residue seems to act as a key not only for the dynamic rearrangement of this intermediate filament protein, but for its role in the integration of other cytoskeletal responses.

## Abbreviations

15d-PGJ2: 15-deoxy-Δ^12,14^-prostaglandin J_2_
DNI: 1,4-dinitro-1H-imidazole
DNP: 2,4-dinitrophenylhydrazone
DNPH: 2,4-dinitrophenylhydrazine
GFP: green fluorescent protein
HNE: 4-hydroxynonenal
HRP: horseradish peroxidase
MEFs: murine embryonic fibroblasts
NI: 4-nitroimidazole
PFA: paraformaldehyde
PTM: Posttranslational modification
RFP: red fluorescent protein
RhoK: Rho kinase
ROI: regions of interest
ROS: reactive oxygen species
TRITC: tetramethylrhodamine B isothiocyanate
ULF: unit length filament
vim: vimentin
wt: wild type.

## Acknowledgements

Feedback from EpiLipidNet (CA19105) and The Summer School organized by SFRRE/FEBS is gratefully acknowledged. We thank the personnel of the Optical Microscopy facility at Centro de Investigaciones Biológicas Margarita Salas for their advice. The technical assistance of MJ Carrasco at the early stages of this work is very much appreciated.

## References

[1] E.M. Hol, Y. Capetanaki, Type III Intermediate Filaments Desmin, Glial Fibrillary Acidic Protein (GFAP), Vimentin, and Peripherin, Cold Spring Harbor perspectives in biology 9(12) (2017).

[2] F. Huber, A. Boire, M.P. Lopez, G.H. Koenderink, Cytoskeletal crosstalk: when three different personalities team up, Curr Opin Cell Biol 32 (2015) 39–47.

[3] M. Muller, S.S. Bhattacharya, T. Moore, Q. Prescott, T. Wedig, H. Herrmann, T.M. Magin, Dominant cataract formation in association with a vimentin assembly disrupting mutation, Hum Mol Genet 18(6) (2009) 1052–7.

[4] B. Cogne, J.E. Bouameur, G. Hayot, X. Latypova, S. Pattabiraman, A. Caillaud, K. Si-Tayeb, T. Besnard, S. Kury, C. Chariau, A. Gaignerie, L. David, P. Bordure, D. Kaganovich, S. Bezieau, C. Golzio, T.M. Magin, B. Isidor, A dominant vimentin variant causes a rare syndrome with premature aging, European journal of human genetics : EJHG (2020).

[5] A. Messing, Alexander disease, Handbook of clinical neurology 148 (2018) 693–700.

[6] Y. Capetanaki, S. Papathanasiou, A. Diokmetzidou, G. Vatsellas, M. Tsikitis, Desmin related disease: a matter of cell survival failure, Curr Opin Cell Biol 32 (2015) 113–20.

[7] R.A. Battaglia, S. Delic, H. Herrmann, N.T. Snider, Vimentin on the move: new developments in cell migration, F1000Research 7 (2018) 1796.

[8] S. Duarte, A. Viedma-Poyatos, E. Navarro-Carrasco, A.E. Martinez, M.A. Pajares, D. Perez-Sala, Vimentin filaments interact with the actin cortex in mitosis allowing normal cell division, Nat Commun 10 (2019) 4200.

[9] G. dos Santos, M.R. Rogel, M.A. Baker, J.R. Troken, D. Urich, L. Morales-Nebreda, J.A. Sennello, M.A. Kutuzov, A. Sitikov, J.M. Davis, A.P. Lam, P. Cheresh, D. Kamp, D.K. Shumaker, G.R. Budinger, K.M. Ridge, Vimentin regulates activation of the NLRP3 inflammasome, Nat Commun 6 (2015) 6574.

[10] I. Ramos, K. Stamatakis, C.L. Oeste, D. Perez-Sala, Vimentin as a Multifaceted Player and Potential Therapeutic Target in Viral Infections, Int J Mol Sci 21(13) (2020) 4675.

[11] K. Strouhalova, M. Prechova, A. Gandalovicova, J. Brabek, M. Gregor, D. Rosel, Vimentin Intermediate Filaments as Potential Target for Cancer Treatment, Cancers 12(1) (2020) 184.

[12] A. Viedma-Poyatos, M.A. Pajares, D. Pérez-Sala, Type III intermediate filaments as targets and effectors of electrophiles and oxidants, Redox Biol 36 (2020) 101582.

[13] Z. Li, J. Wu, J. Zhou, B. Yuan, J. Chen, W. Wu, L. Mo, Z. Qu, F. Zhou, Y. Dong, K. Huang, Z. Liu, T. Wang, D. Symmes, J. Gu, E. Sho, J. Zhang, R. Chen, Y. Xu, A Vimentin-Targeting Oral Compound with Host-Directed Antiviral and Anti-Inflammatory Actions Addresses Multiple Features of COVID-19 and Related Diseases, mBio 12(5) (2021) e0254221.

[14] K.M. Ridge, J.E. Eriksson, M. Pekny, R.D. Goldman, Roles of vimentin in health and disease, Genes Dev 36(7-8) (2022) 391–407.

[15] H.J. Forman, H. Zhang, Targeting oxidative stress in disease: promise and limitations of antioxidant therapy, Nat Rev Drug Discov 20(9) (2021) 689–709.

[16] D. Pérez-Sala, C.L. Oeste, A.E. Martínez, B. Garzón, M.J. Carrasco, F.J. Cañada, Vimentin filament organization and stress sensing depend on its single cysteine residue and zinc binding, Nat Commun 6 (2015) 7287.

[17] S. Duarte, T. Melo, R. Domingues, J.d.D. Alché, D. Pérez-Sala, Insight into the cellular effects of nitrated phospholipids: evidence for pleiotropic mechanisms of action, Free Rad Biol Med 144 (2019) 192–202.

[18] A. Mónico, S. Duarte, M.A. Pajares, D. Pérez-Sala, Vimentin disruption by lipoxidation and electrophiles: role of the cysteine residue and filament dynamics, Redox Biol 23 (2019) 101098.

[19] A. Viedma-Poyatos, P. Gonzalez-Jimenez, O. Langlois, I. Company-Marin, C.M. Spickett, D. Perez-Sala, Protein Lipoxidation: Basic Concepts and Emerging Roles, Antioxidants (Basel) 10(2) (2021) e1009128.

[20] H. Herrmann, U. Aebi, Intermediate filaments: molecular structure, assembly mechanism, and integration into functionally distinct intracellular Scaffolds, Annu Rev Biochem 73 (2004) 749–89.

[21] A. Premchandar, N. Mucke, J. Poznanski, T. Wedig, M. Kaus-Drobek, H. Herrmann, M. Dadlez, Structural Dynamics of the Vimentin Coiled-coil Contact Regions Involved in Filament Assembly as Revealed by Hydrogen-Deuterium Exchange, J Biol Chem 291(48) (2016) 24931–24950.

[22] J.F. Hess, M.S. Budamagunta, A. Aziz, P.G. FitzGerald, J.C. Voss, Electron paramagnetic resonance analysis of the vimentin tail domain reveals points of order in a largely disordered region and conformational adaptation upon filament assembly, Protein science : a publication of the Protein Society 22(1) (2013) 47–55.

[23] J.T. Pittenger, J.F. Hess, M.S. Budamagunta, J.C. Voss, P.G. Fitzgerald, Identification of phosphorylation-induced changes in vimentin intermediate filaments by site-directed spin labeling and electron paramagnetic resonance, Biochemistry 47(41) (2008) 10863–70.

[24] E. Griesser, V. Vemula, A. Mónico, D. Pérez-Sala, M. Fedorova, Dynamic posttranslational modifications of cytoskeletal proteins unveil hot spots under nitroxidative stress, Redox Biol 44 (2021) 102014.

[25] D. Frescas, C.M. Roux, S. Aygun-Sunar, A.S. Gleiberman, P. Krasnov, O.V. Kurnasov, E. Strom, L.P. Virtuoso, M. Wrobel, A.L. Osterman, M.P. Antoch, V. Mett, O.B. Chernova, A.V. Gudkov, Senescent cells expose and secrete an oxidized form of membrane-bound vimentin as revealed by a natural polyreactive antibody, Proc Natl Acad Sci U S A 114(9) (2017) E1668–E1677.

[26] T. Uemura, T. Suzuki, K. Ko, M. Nakamura, N. Dohmae, A. Sakamoto, Y. Terui, T. Toida, K. Kashiwagi, K. Igarashi, Structural change and degradation of cytoskeleton due to the acrolein conjugation with vimentin and actin during brain infarction, Cytoskeleton (Hoboken) 77 (2020) 414–421.

[27] I.M. Riederer, R.M. Herrero, G. Leuba, B.M. Riederer, Serial protein labeling with infrared maleimide dyes to identify cysteine modifications, J Proteomics 71(2) (2008) 222–30.

[28] A. Mónico, J. Guzman-Caldentey, M.A. Pajares, S. Martin-Santamaria, D. Pérez-Sala, Molecular Insight into the Regulation of Vimentin by Cysteine Modifications and Zinc Binding, Antioxidants (Basel) 10(7) (2021) 1039.

[29] K.R. Rogers, H. Herrmann, W.W. Franke, Characterization of disulfide crosslink formation of human vimentin at the dimer, tetramer, and intermediate filament levels, J Struct Biol 117(1) (1996) 55–69.

[30] S. Duarte, A. Viedma-Poyatos, A. Mónico, D. Pérez-Sala, The conserved cysteine residue of type III intermediate filaments serves as a structural element and redox sensor, Free Rad Biol Med 120 (2018) S84.

[31] M. Kaus-Drobek, N. Mucke, R.H. Szczepanowski, T. Wedig, M. Czarnocki-Cieciura, M. Polakowska, H. Herrmann, A. Wyslouch-Cieszynska, M. Dadlez, Vimentin S-glutathionylation at Cys328 inhibits filament elongation and induces severing of mature filaments in vitro, FEBS J 287 (2020) 5304–5322.

[32] M. Fratelli, H. Demol, M. Puype, S. Casagrande, I. Eberini, M. Salmona, V. Bonetto, M. Mengozzi, F. Duffieux, E. Miclet, A. Bachi, J. Vandekerckhove, E. Gianazza, P. Ghezzi, Identification by redox proteomics of glutathionylated proteins in oxidatively stressed human T lymphocytes, Proc Natl Acad Sci U S A 99 (2002) 3505–3510.

[33] M.B. West, B.G. Hill, Y.T. Xuan, A. Bhatnagar, Protein glutathiolation by nitric oxide: an intracellular mechanism regulating redox protein modification, Faseb J 20(10) (2006) 1715–7.

[34] K. Stamatakis, F.J. Sánchez-Gómez, D. Pérez-Sala, Identification of novel protein targets for modification by 15-deoxy-D^12,14^-prostaglandin J_2_ in mesangial cells reveals multiple interactions with the cytoskeleton., J Am Soc Nephrol 17 (2006) 89–98.

[35] J. Chavez, W.G. Chung, C.L. Miranda, M. Singhal, J.F. Stevens, C.S. Maier, Site-specific protein adducts of 4-hydroxy-2(E)-nonenal in human THP-1 monocytic cells: protein carbonylation is diminished by ascorbic acid, Chem Res Toxicol 23(1) (2010) 37–47.

[36] O. Hernández-Perera, D. Pérez-Sala, E. Soria, S. Lamas, Involvement of Rho GTPases in the transcriptional inhibition of pre-proendothelin-1 gene expression by Simvastatin in vascular endothelial cells, Circ. Res. 87 (2000) 616–622.

[37] C. Kruger, S.J. Burke, J.J. Collier, T.T. Nguyen, J.M. Salbaum, K. Stadler, Lipid peroxidation regulates podocyte migration and cytoskeletal structure through redox sensitive RhoA signaling, Redox Biol 16 (2018) 248–254.

[38] A.J. Sarria, S.K. Nordeen, R.M. Evans, Regulated expression of vimentin cDNA in cells in the presence and absence of a preexisting vimentin filament network, J Cell Biol 111(2) (1990) 553–65.

[39] A.J. Sarria, J.G. Lieber, S.K. Nordeen, R.M. Evans, The presence or absence of a vimentin-type intermediate filament network affects the shape of the nucleus in human SW-13 cells, J Cell Sci 107 (Pt 6) (1994) 1593–607.

[40] C.L. Oeste, D. Pérez-Sala, Modification of cysteine residues by cyclopentenone prostaglandins: interplay with redox regulation of protein function, Mass spectrometry reviews 33 (2014) 110–125.

[41] S.Y. Shin, J. Park, Y. Jung, Y.H. Lee, D. Koh, Y. Yoon, Y. Lim, Anticancer activities of cyclohexenone derivatives, Appl Biol Chem 63 (2020) 82.

[42] I. Cervenka, L.Z. Agudelo, J.L. Ruas, Kynurenines: Tryptophan’s metabolites in exercise, inflammation, and mental health, Science 357(6349) (2017).

[43] H.Q. Wu, P. Guidetti, J.H. Goodman, M. Varasi, G. Ceresoli-Borroni, C. Speciale, H.E. Scharfman, R. Schwarcz, Kynurenergic manipulations influence excitatory synaptic function and excitotoxic vulnerability in the rat hippocampus in vivo, Neuroscience 97(2) (2000) 243–51.

[44] L. Gallego-Villar, L. Hannibal, J. Haberle, B. Thony, T. Ben-Omran, G.K. Nasrallah, A.N. Dewik, W.D. Kruger, H.J. Blom, Cysteamine revisited: repair of arginine to cysteine mutations, Journal of inherited metabolic disease 40(4) (2017) 555–567.

[45] D. Leitsch, D. Kolarich, I.B. Wilson, F. Altmann, M. Duchene, Nitroimidazole action in Entamoeba histolytica: a central role for thioredoxin reductase, PLoS Biol 5(8) (2007) e211.

[46] C.L. Ho, J.L. Martys, A. Mikhailov, G.G. Gundersen, R.K. Liem, Novel features of intermediate filament dynamics revealed by green fluorescent protein chimeras, J Cell Sci 111 (Pt 13) (1998) 1767–78.

[47] F.M. Notarangelo, X.D. Wang, K.J. Horning, R. Schwarcz, Role of d-amino acid oxidase in the production of kynurenine pathway metabolites from d-tryptophan in mice, J Neurochem 136(4) (2016) 804–814.

[48] Q. Luo, Y. Tao, W. Sheng, J. Lu, H. Wang, Dinitroimidazoles as bifunctional bioconjugation reagents for protein functionalization and peptide macrocyclization, Nat Commun 10(1) (2019) 142.

[49] H. Esterbauer, R.J. Schaur, H. Zollner, Chemistry and biochemistry of 4-hydroxynonenal, malonaldehyde and related aldehydes, Free Radic Biol Med 11(1) (1991) 81–128.

[50] S. Vazquez, J.A. Aquilina, J.F. Jamie, M.M. Sheil, R.J. Truscott, Novel protein modification by kynurenine in human lenses, J Biol Chem 277(7) (2002) 4867–73.

[51] N.R. Parker, A. Korlimbinis, J.F. Jamie, M.J. Davies, R.J. Truscott, Reversible binding of kynurenine to lens proteins: potential protection by glutathione in young lenses, Invest Ophthalmol Vis Sci 48(8) (2007) 3705–13.

[52] K. Uchida, E.R. Stadtman, Modification of histidine residues in proteins by reaction with 4-hydroxynonenal, Proc Natl Acad Sci U S A 89(10) (1992) 4544–8.

[53] H. Song, H. Park, Y.S. Kim, K.D. Kim, H.K. Lee, D.H. Cho, J.W. Yang, D.Y. Hur, L-kynurenine-induced apoptosis in human NK cells is mediated by reactive oxygen species, International immunopharmacology 11(8) (2011) 932–8.

[54] A. Sannella, L. Gradoni, T. Persichini, E. Ongini, G. Venturini, M. Colasanti, Intracellular release of nitric oxide by NCX 972, an NO-releasing metronidazole, enhances in vitro killing of Entamoeba histolytica, Antimicrob Agents Chemother 47(7) (2003) 2303–6.

[55] A.M. Rice, Y. Long, S.B. King, Nitroaromatic Antibiotics as Nitrogen Oxide Sources, Biomolecules 11(2) (2021) 267.

[56] M.A. Mackinder, C.A. Evans, J. Chowdry, C.A. Staton, B.M. Corfe, Alteration in composition of keratin intermediate filaments in a model of breast cancer progression and the potential to reverse hallmarks of metastasis, Cancer biomarkers : section A of Disease markers 12(2) (2012) 49–64.

[57] G. Aldini, M.R. Domingues, C.M. Spickett, P. Domingues, A. Altomare, F.J. Sánchez-Gómez, C.L. Oeste, D. Pérez-Sala, Protein lipoxidation: detection strategies and challenges, Redox Biol 5 (2015) 253–266.

[58] I. Lois-Bermejo, P. González-Jiménez, S. Duarte, M.A. Pajares, D. Pérez-Sala, Vimentin tail segments are differentially exposed at distinct cellular locations and in response to stress, Frontiers in cell and developmental biology 10 (2022) 908263.

[59] I. Dalle-Donne, R. Rossi, A. Milzani, P. Di Simplicio, R. Colombo, The actin cytoskeleton response to oxidants: from small heat shock protein phosphorylation to changes in the redox state of actin itself, Free Radic Biol Med 31(12) (2001) 1624–32.

[60] C. Wilson, J.R. Terman, C. Gonzalez-Billault, G. Ahmed, Actin filaments-A target for redox regulation, Cytoskeleton (Hoboken) 73(10) (2016) 577–595.

[61] S. Varland, J. Vandekerckhove, A. Drazic, Actin Post-translational Modifications: The Cinderella of Cytoskeletal Control, Trends Biochem Sci 44(6) (2019) 502–516.

[62] E.K. Ahmed, A. Rogowska-Wrzesinska, P. Roepstorff, A.L. Bulteau, B. Friguet, Protein modification and replicative senescence of WI-38 human embryonic fibroblasts, Aging cell 9(2) (2010) 252–72.

[63] A. Swiader, C. Camare, P. Guerby, R. Salvayre, A. Negre-Salvayre, 4-Hydroxynonenal Contributes to Fibroblast Senescence in Skin Photoaging Evoked by UV-A Radiation, Antioxidants (Basel) 10(3) (2021) 365.

[64] Y. Miyata, J.N. Rauch, U.K. Jinwal, A.D. Thompson, S. Srinivasan, C.A. Dickey, J.E. Gestwicki, Cysteine reactivity distinguishes redox sensing by the heat-inducible and constitutive forms of heat shock protein 70, Chemistry & biology 19(11) (2012) 1391–9.

[65] S.E. Permyakov, E.Y. Zernii, E.L. Knyazeva, A.I. Denesyuk, A.A. Nazipova, T.V. Kolpakova, D.V. Zinchenko, P.P. Philippov, E.A. Permyakov, Senin, II, Oxidation mimicking substitution of conservative cysteine in recoverin suppresses its membrane association, Amino acids 42(4) (2012) 1435–42.

[66] E. Amin, B.N. Dubey, S.C. Zhang, L. Gremer, R. Dvorsky, J.M. Moll, M.S. Taha, L. Nagel-Steger, R.P. Piekorz, A.V. Somlyo, M.R. Ahmadian, Rho-kinase: regulation, (dys)function, and inhibition, Biol Chem 394(11) (2013) 1399–410.

[67] S. Hartmann, A.J. Ridley, S. Lutz, The Function of Rho-Associated Kinases ROCK1 and ROCK2 in the Pathogenesis of Cardiovascular Disease, Frontiers in pharmacology 6 (2015) 276.

[68] Y. Jiu, J. Peranen, N. Schaible, F. Cheng, J.E. Eriksson, R. Krishnan, P. Lappalainen, Vimentin intermediate filaments control actin stress fiber assembly through GEF-H1 and RhoA, J Cell Sci 130(5) (2017) 892–902.

[69] S.J. Kim, Z.W. Lee, S.M. Kweon, S. Kim, K.S. Ha, Regulation of reactive oxygen species and stress fiber formation by calpeptin in Swiss 3T3 fibroblasts, Cellular signalling 14(3) (2002) 205–10.

[70] A.L. Berr, K. Wiese, G. dos Santos, J.M. Davis, C.M. Koch, K.R. Anekalla, M.E. Kidd, Y. Cheng, Y.-S. Hu, K.M. Ridge, Vimentin is required for tumor progression and metastasis in a mouse model of non-small cell lung cancer, BioRxiv (2020) doi: https://doi.org/10.1101/2020.06.04.130963.

[71] S. Sivagurunathan, A. Vahabikashi, H. Yang, J. Zhang, K. Vazquez, D. Rajasundaram, Y. Politanska, H. Abdala-Valencia, J. Notbohm, M. Guo, S.A. Adam, R.D. Goldman, Expression of vimentin alters cell mechanics, cell-cell adhesion, and gene expression profiles suggesting the induction of a hybrid EMT in human mammary epithelial cells, Frontiers in cell and developmental biology 10 (2022) 929495.

[72] G. Aldini, M. Carini, G. Vistoli, T. Shibata, Y. Kusano, L. Gamberoni, I. Dalle-Donne, A. Milzani, K. Uchida, Identification of Actin as a 15-Deoxy-Delta(12,14)-prostaglandin J(2) Target in Neuroblastoma Cells: Mass Spectrometric, Computational, and Functional Approaches To Investigate the Effect on Cytoskeletal Derangement, Biochemistry 46(10) (2007) 2707–2718.

[73] L.M. Landino, M.T. Koumas, C.E. Mason, J.A. Alston, Modification of tubulin cysteines by nitric oxide and nitroxyl donors alters tubulin polymerization activity, Chem Res Toxicol 20(11) (2007) 1693–700.

[74] V. Pekovic, I. Gibbs-Seymour, E. Markiewicz, F. Alzoghaibi, A.M. Benham, R. Edwards, M. Wenhert, T. von Zglinicki, C.J. Hutchison, Conserved cysteine residues in the mammalian lamin A tail are essential for cellular responses to ROS generation, Aging cell 10(6) (2011) 1067–79.

[75] S. Gharbi, B. Garzón, J. Gayarre, J. Timms, D. Pérez-Sala, Study of protein targets for covalent modification by the antitumoral and anti-inflammatory prostaglandin PGA_1_: focus on vimentin, J Mass Spectrom 42 (2007) 1474–1484.

[76] R.M. LoPachin, T. Gavin, D.R. Petersen, D.S. Barber, Molecular mechanisms of 4-hydroxy-2-nonenal and acrolein toxicity: nucleophilic targets and adduct formation, Chem Res Toxicol 22(9) (2009) 1499–508.

[77] M.J. Davies, Protein oxidation and peroxidation, Biochem J 473(7) (2016) 805–25.

[78] M.J. Davies, The oxidative environment and protein damage, Biochim Biophys Acta 1703(2) (2005) 93–109.

[79] K. Uchida, S. Kawakishi, Identification of oxidized histidine generated at the active site of Cu,Zn-superoxide dismutase exposed to H2O2. Selective generation of 2-oxo-histidine at the histidine 118, J Biol Chem 269(4) (1994) 2405–10.

[80] P.V. Usatyuk, V. Natarajan, Role of Mitogen-activated Protein Kinases in 4-Hydroxy-2-nonenalinduced Actin Remodeling and Barrier Function in Endothelial Cells, J Biol Chem 279 (2004) 11789–11797.

[81] H. Sies, D.P. Jones, Reactive oxygen species (ROS) as pleiotropic physiological signalling agents, Nat Rev Mol Cell Biol 21(7) (2020) 363–383.

[82] J.M. Chalker, G.J. Bernardes, Y.A. Lin, B.G. Davis, Chemical modification of proteins at cysteine: opportunities in chemistry and biology, Chemistry, an Asian journal 4(5) (2009) 630–40.

[83] L.S. Havel, E.R. Kline, A.M. Salgueiro, A.I. Marcus, Vimentin regulates lung cancer cell adhesion through a VAV2-Rac1 pathway to control focal adhesion kinase activity, Oncogene 34(15) (2015) 1979–90.

[84] J. Huot, F. Houle, F. Marceau, J. Landry, Oxidative stress-induced actin reorganization mediated by the p38 mitogen-activated protein kinase/heat shock protein 27 pathway in vascular endothelial cells, Circ Res 80(3) (1997) 383–92.

[85] Y. Zhao, H.W. Davis, Hydrogen peroxide-induced cytoskeletal rearrangement in cultured pulmonary endothelial cells, J Cell Physiol 174(3) (1998) 370–9.

[86] M. Gregor, S. Osmanagic-Myers, G. Burgstaller, M. Wolfram, I. Fischer, G. Walko, G.P. Resch, A. Jorgl, H. Herrmann, G. Wiche, Mechanosensing through focal adhesion-anchored intermediate filaments, FASEB J 28(2) (2014) 715–29.

[87] A. Aghajanian, E.S. Wittchen, S.L. Campbell, K. Burridge, Direct activation of RhoA by reactive oxygen species requires a redox-sensitive motif, PLoS One 4(11) (2009) e8045.

[88] J.S. Kim, J.G. Kim, C.Y. Jeon, H.Y. Won, M.Y. Moon, J.Y. Seo, J.I. Kim, J. Kim, J.Y. Lee, S.Y. Choi, J. Park, J.H. Yoon Park, K.S. Ha, P.H. Kim, J.B. Park, Downstream components of RhoA required for signal pathway of superoxide formation during phagocytosis of serum opsonized zymosans in macrophages, Experimental & molecular medicine 37(6) (2005) 575–87.

[89] H. Goto, H. Kosako, K. Tanabe, M. Yanagida, M. Sakurai, M. Amano, K. Kaibuchi, M. Inagaki, Phosphorylation of vimentin by Rho-associated kinase at a unique amino-terminal site that is specifically phosphorylated during cytokinesis, J Biol Chem 273(19) (1998) 11728–36.

[90] R. Gerhard, H. John, K. Aktories, I. Just, Thiol-modifying phenylarsine oxide inhibits guanine nucleotide binding of Rho but not of Rac GTPases, Mol Pharmacol 63(6) (2003) 1349–55.

[91] H. Wu, Y. Shen, S. Sivagurunathan, M.S. Weber, S.A. Adam, J.H. Shin, J.J. Fredberg, O. Medalia, R. Goldman, D.A. Weitz, Vimentin intermediate filaments and filamentous actin form unexpected interpenetrating networks that redefine the cell cortex, Proc Natl Acad Sci U S A 119(10) (2022) e2115217119.

[92] I. Ding, Z. Ostrowska-Podhorodecka, W. Lee, R.S.C. Liu, K. Carneiro, P.A. Janmey, C.A. McCulloch, Cooperative roles of PAK1 and filamin A in regulation of vimentin assembly and cell extension formation, Biochim Biophys Acta Mol Cell Res 1867(9) (2020) 118739.

[93] G. Wiche, Plectin-Mediated Intermediate Filament Functions: Why Isoforms Matter, Cells 10(8) (2021) 2154.

[94] R. Spurny, K. Abdoulrahman, L. Janda, D. Runzler, G. Kohler, M.J. Castanon, G. Wiche, Oxidation and Nitrosylation of Cysteines Proximal to the Intermediate Filament (IF)-binding Site of Plectin: effects on structure and vimentin binding and involvement in if collapse, J Biol Chem 282(11) (2007) 8175–87.

[95] K.T. Pan, Y.Y. Chen, T.H. Pu, Y.S. Chao, C.Y. Yang, R.D. Bomgarden, J.C. Rogers, T.C. Meng, K.H. Khoo, Mass spectrometry-based quantitative proteomics for dissecting multiplexed redox cysteine modifications in nitric oxide-protected cardiomyocyte under hypoxia, Antioxid Redox Signal 20(9) (2014) 1365–81.

