## Supplementary material for "Vimentin cysteine 328 modifications finely tune network organization and influence actin remodeling under oxidative and electrophilic stress": Suppl.

Running title: Role of vimentin C328 in cytoskeletal interplay

Supplementary Information

Supplementary Table 1

| Vimentin | Template | Sequence 5'-3' |
|---|---|---|
| wt | - | GGTGCAGTCCCTCACCTGTGAAGTGGATGCCC |
| C328S | Vimentin wt | GGTGCAGTCCCTCACC**TCT**GAAGTGGATGCCC |
| C328A | Vimentin wt | GGTGCAGTCCCTCACC**GCT**GAAGTGGATGCCC |
| C328H | Vimentin wt | GGTGCAGTCCCTCACC**CAT**GAAGTGGATGCCC |
| C328F | Vimentin wt | GGTGCAGTCCCTCACC**TTT**GAAGTGGATGCCC |
| C328W | Vimentin wt | GGTGCAGTCCCTCACC**TGG**GAAGTGGATGCCC |
| C328D | Vimentin C328H | GGTGCAGTCCCTCACC**GAT**GAAGTGGATGCCC |

The mutated codon is shown in bold.

Supplementary Information

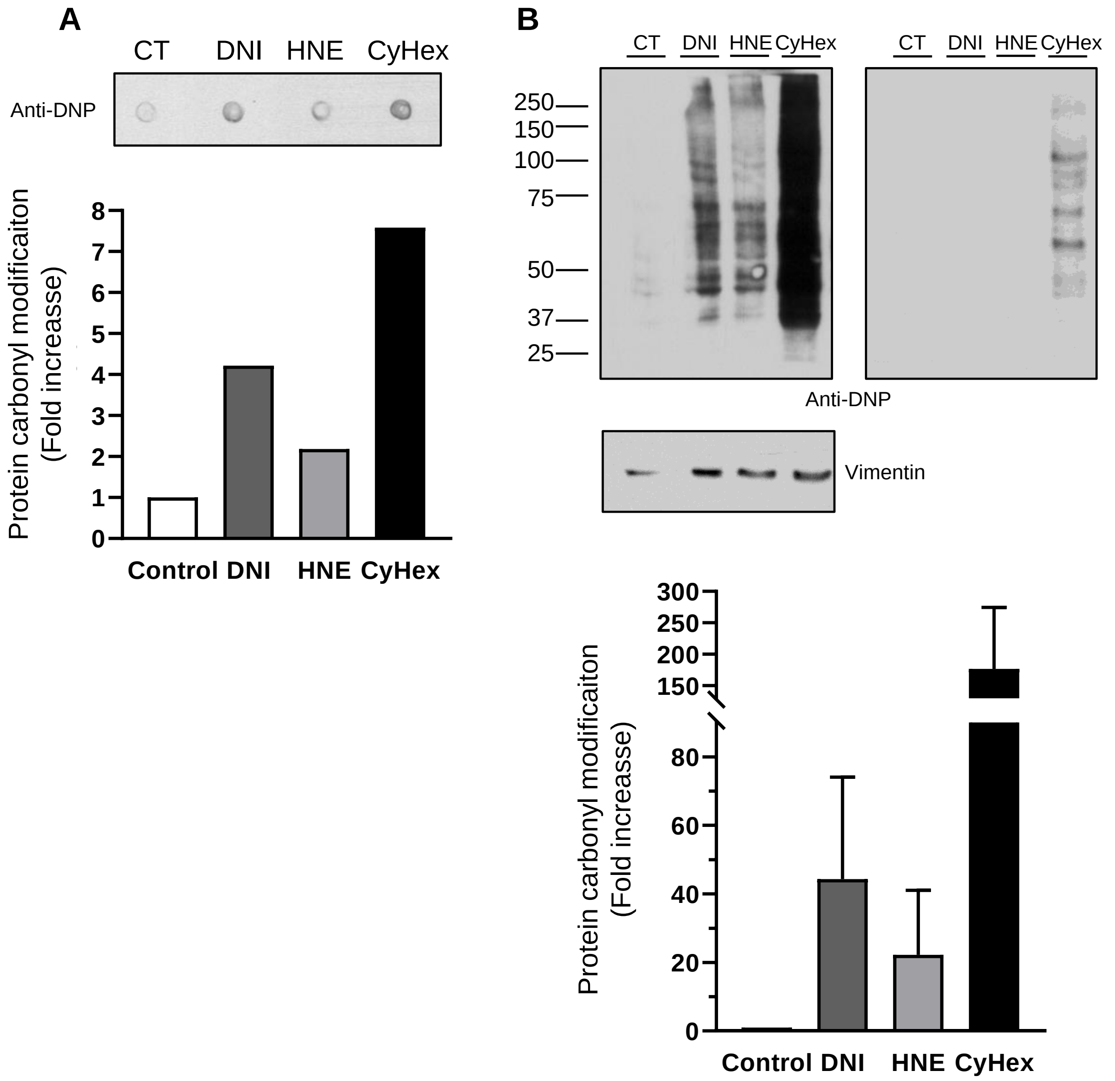


**Supplementary Figure 1. Effect of treatment of cells with various electrophiles on protein carbonyl formation.** SW13/cl.2 cells (A) or SW13/cl.2 cells expressing GFP-vimentin wt (B) were treated with the indicated compounds as described above. Cell lysates were subjected to derivatization of protein carbonyls using OxyBlot Protein Oxidation Detection Kit, and aliquots were spotted on membrane (A) or separated by SDS-PAGE and transferred to Immobilon membranes (B) for detection of protein carbonyls with the anti-DNP antibody. Two different exposures of the same membrane incubated with anti-DNP are shown in panel B. (A) Results are representative of two assays. (B) Results are representative from at least five Oxyblot experiments. The position of the molecular weight standards is shown on the left of the blots. Quantitation of protein carbonyl modifications are shown in the corresponding graphs as values relatives to the signal of the control condition. CyHex, cyclohexenone.

Supplementary Information

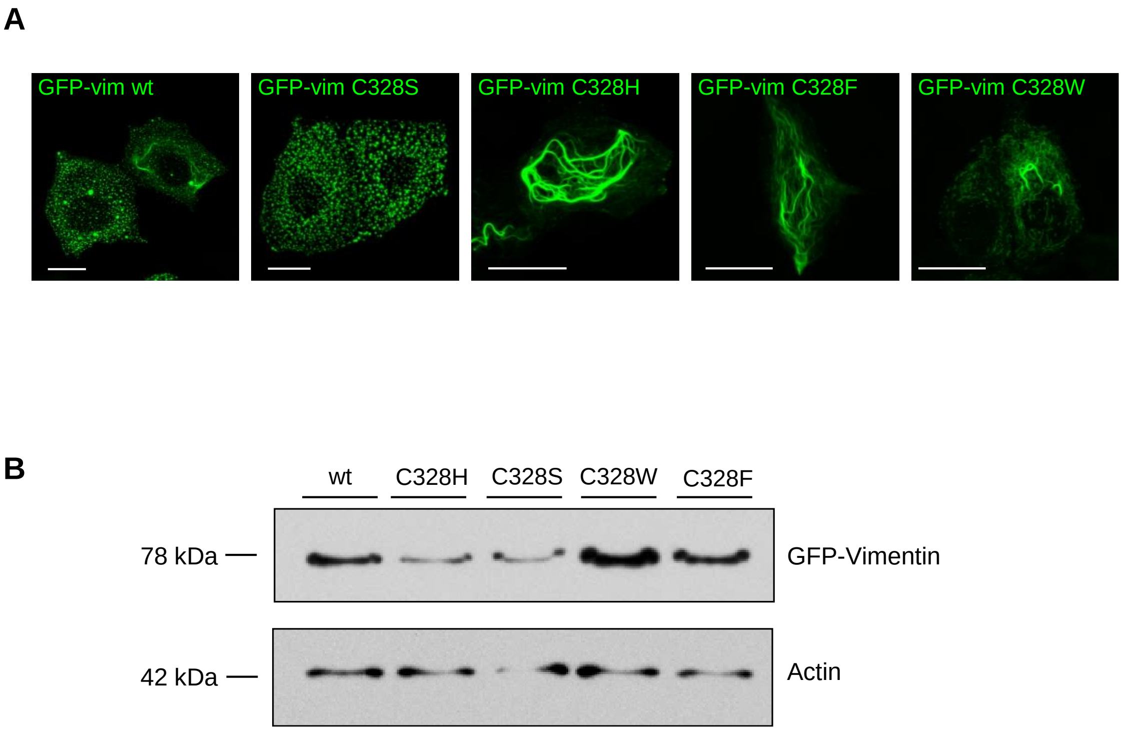


**Supplementary Figure 2. Expression of GFP-vimentin wt and various cysteine mutants in vimentin-deficient MCF7 cells.** MCF7 cells were transiently transfected with the indicated constructs. (A) The morphology of vimentin assemblies was monitored by fluorescence microscopy. Overall projections, representative from three different experiments are shown. Bars, 20 µm. (B) Lysates from MCF7 cells transfected with the indicated constructs were analyzed by SDS-PAGE and western blot with anti-vimentin antibody. Levels of actin were used as loading control.

Supplementary Information

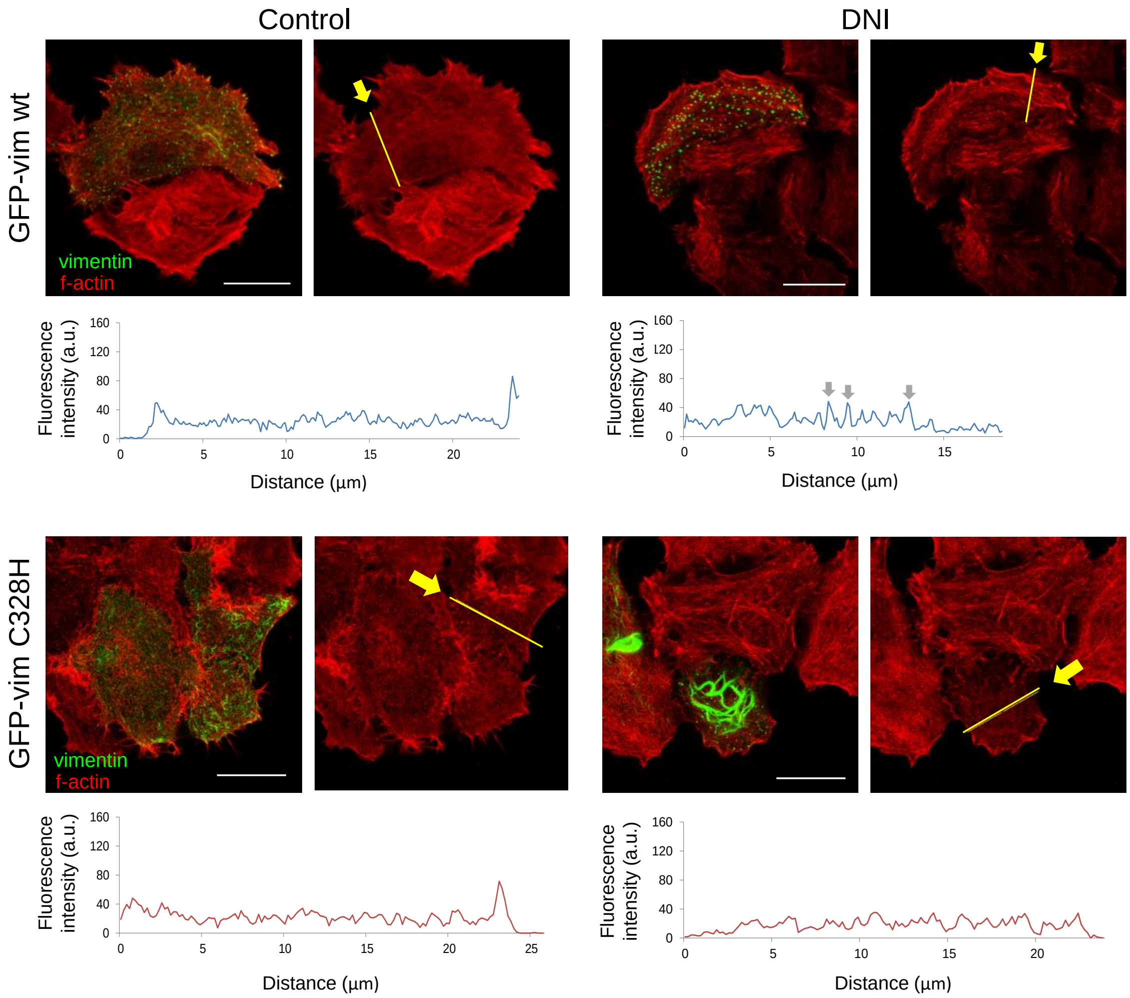


**Supplementary Figure 3. Effect of DNI on actin remodeling in MCF7 cells expressing a GFP-vimentin wt or C328H.** MCF7 cells transiently transfected with GFP-vimentin wt or C328H were treated with DNI, as indicated. After treatment, cells were fixed, f-actin was stained with phalloidin-TRITC and the distribution of GFP-vimentin (green) and f-actin (red) was assessed by confocal microscopy. Single sections were taken from the lower third of cells and merged channels (left) and f-actin staining (right) are shown. Fluorescence intensity profiles of f-actin distribution were obtained along the lines drawn on images in the direction indicated by arrows. The position of stress fibers is indicated on the profiles by grey arrows.

Supplementary Information

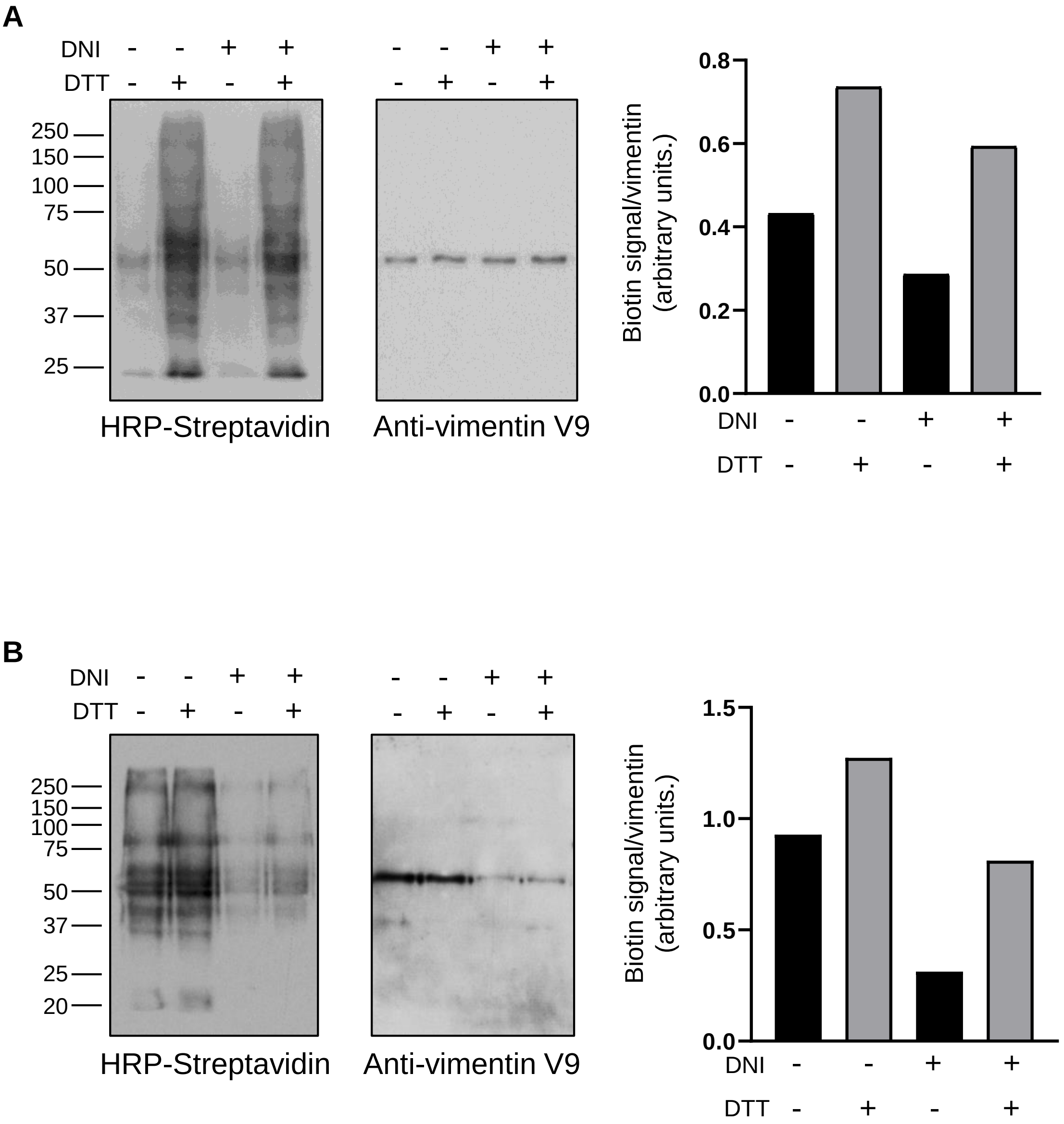


**Supplementary Figure 4. Effect of DNI on cysteine accessibility in cells expressing vimentin wt.** SW13/cl.2 cells stably transfected with RFP//vimentin wt were treated with DNI, as indicated. Accessibility of cysteine residues in vimentin was evaluated by incubation of total cell lysates from control or DNI-treated cells in the absence or presence of DTT, and subsequent incubation with biotinylated iodoacetamide (A) or biotinylated maleimide (B), followed by vimentin immunoprecipitation and assessment of the incorporation of the corresponding biotinylated reagents in immunoprecipitates by SDS-PAGE, blot and biotin detection with HRP-streptavidin. Anti-vimentin V9 antibody was used to estimate the amount of vimentin present in the immunoprecipitation. Results shown in each case are representative from two assays performed in duplicate.
